# PL25_M23: Salt-enhanced antibacterial activity of a plasmid-encoded SLT–M23 hydrolase from *Virgibacillus salarius* against Gram-positive bacteria

**DOI:** 10.64898/2026.11.24.768938

**Authors:** Ayman H. Farran, Nicholas Kontis, Alexandre S. Rosado

## Abstract

Antimicrobial resistance has intensified interest in peptidoglycan hydrolases as potential alternatives to conventional antibiotics. However, halophilic and halotolerant bacteria remain underexplored sources of these enzymes. Here, we characterized PL25_M23, a predicted 370-amino-acid, dual-domain cell wall hydrolase encoded on a candidate closed circular plasmid from *Virgibacillus salarius* PL25, a halotolerant Gram-positive isolate. PL25_M23 combines an N-terminal Slt35/lysozyme-like glycoside hydrolase fold with a C-terminal Peptidase_M23 domain containing the canonical HxxxD and HxH Zn^2+^-binding motifs characteristic of the M23 family. Although its closest characterized homolog, CwlP, shares limited sequence identity with PL25_M23, AlphaFold2 modeling and domain-restricted structural superposition support a CwlP-like, two-domain architecture. The predicted protein surface is enriched in acidic residues, a compositional feature commonly reported in halophilic proteins. Recombinant PL25_M23 reduced the viable counts of seven Gram-positive strains representing four genera, including methicillin-sensitive and methicillin-resistant *Staphylococcus aureus*, by approximately 1.9–3.8 log_10_ CFU/mL at a concentration of 100 µg/mL within 60 min. EDTA treatment reduced PL25_M23 activity, whereas Zn^2+^ supplementation restored antibacterial activity, resulting in reductions of 1.4–3.6 log_10_ CFU/mL and indicating that its antibacterial activity is sensitive to metal chelation. The extent of bacterial killing increased with NaCl supplementation in all salt-tolerant test strains, with an increase of 1.4 log_10_ CFU/mL. Together, these findings identify PL25_M23 as a divergent, plasmid-encoded hydrolase with antibacterial activity against the Gram-positive strains tested and establish it as a candidate for further biochemical characterization and evaluation as an enzybiotic.

## 1. Introduction

Antimicrobial resistance is increasing the burden of infections that are difficult to treat with conventional antibiotics (Murray et al., 2022). Among Gram-positive pathogens, methicillin-resistant *Staphylococcus aureus* (MRSA) and vancomycin-resistant *Enterococcus faecium* remain high-priority targets for the development of novel antibacterial strategies (Sati et al., 2025). Peptidoglycan (PG) hydrolases are promising candidates for enzybiotic applications because they directly cleave bonds within the bacterial cell wall, thereby inducing rapid lysis of target cells (Vollmer, 2008; Liu et al., 2023; Sabur et al., 2025). Among these enzymes, members of the Peptidase_M23 family include well-characterized metallopeptidases, many of which are Zn^2+^-dependent DD-endopeptidases that cleave specific peptide cross-links within PG (Razew et al., 2022). A prototypical M23 enzyme, lysostaphin, is produced by *Staphylococcus simulans* biovar *staphylolyticus* and cleaves the pentaglycine cross-bridges in the PG of *S. aureus* (Bastos et al., 2010; Charoenjotivadhanakul et al., 2023). Other members of this family include LytM from *S. aureus* (Firczuk et al., 2005; Antenucci et al., 2024), ALE-1 from *Staphylococcus capitis* (Sugai et al., 1997), zoocin A from *Streptococcus equi* subsp. *zooepidemicus* (Simmonds et al., 1996), and EnpA from *Enterococcus faecalis* prophage (Małecki et al., 2021). Although these enzymes differ in substrate specificity and biological context, characterized Peptidase_M23 enzymes commonly contain conserved HxxxD and HxH motifs that constitute the canonical metal-binding architecture of their catalytic domains (Firczuk et al., 2005; Razew et al., 2022). Mobile genetic elements can also encode multidomain PG hydrolases that combine complementary catalytic modules within a single protein (Razew et al., 2022). For example, CwlT, which is encoded by the *Bacillus subtilis* integrative and conjugative element ICEBs1, contains an N-terminal GH23-like bacterial lysozyme domain and a C-terminal NlpC/P60 peptidase domain. Together, these domains contribute to the remodeling of donor-cell PG during element transfer (DeWitt and Grossman, 2014). Another example is CwlP (YomI), which is encoded by the SP-β prophage of *B. subtilis* and combines an N-terminal lytic transglycosylase (SLT) domain with a C-terminal M23 peptidase domain (Sudiarta et al., 2010). Interestingly, the M23-dependent activity of CwlP has been reported to be independent of Zn^2+^, despite the general association of M23-family enzymes with Zn^2+^-dependent catalysis (Sudiarta et al., 2010; Razew et al., 2022). These examples illustrate how mobile genetic elements can encode multidomain hydrolases that combine complementary lytic activities for specialized cell-wall remodeling (Sudiarta et al., 2010; DeWitt and Grossman, 2014).

Despite the growing number of characterized PG hydrolases, enzymes derived from halophilic and halotolerant members of the Bacillaceae family remain poorly characterized (Shockman et al., 1993; Ventosa et al., 1998; Van Heijenoort, 2011). Halophilic proteins are typically enriched in acidic surface residues and have relatively low isoelectric points, physicochemical characteristics associated with protein stability and solubility under high-ionic-strength conditions (Madern et al., 2000; Graziano and Merlino, 2014; Kougentakis et al., 2020). In addition, halophilic and halotolerant microorganisms are recognized as valuable sources of hydrolytic enzymes with distinctive biochemical properties and the ability to function under high-salinity conditions (De Lourdes Moreno et al., 2013; Amoozegar and Siroosi, 2015). However, plasmid-associated PG hydrolases from these microorganisms remain largely unexplored.

We hypothesized that PL25_M23, a cell wall hydrolase encoded by a candidate plasmid from the halotolerant bacterium *Virgibacillus salarius* PL25, could reduce viable counts of Gram-positive bacteria while retaining or exhibiting enhanced activity under high-salinity conditions. To investigate this possibility, we characterized pPL25-1, a 74-kb closed circular plasmid candidate with unverified mobility, from *V. salarius* PL25 and examined its encoded 370-amino-acid predicted dual-domain SLT/M23 hydrolase using genomic, phylogenetic, structural, and functional analyses. We subsequently evaluated the antibacterial activity of recombinant PL25_M23 against seven Gram-positive strains, including methicillin-resistant and methicillin-sensitive *S. aureus* (MRSA and MSSA, respectively), and investigated the effects of elevated salinity on its activity.

## 2. Materials and Methods

Genome-based taxonomic assignment was performed using the Type Strain Genome Server (TYGS) (Meier-Kolthoff and Göker, 2019). Digital DNA–DNA hybridization (dDDH) values were calculated against type-strain genomes using the Genome-to-Genome Distance Calculator (GGDC) formula d_4_, with a threshold of ≥70% used for species delineation. TYGS also generated a whole-genome phylogeny by comparing the query genome with genomes in its type-strain database. Branch support was assessed using 100 pseudo-bootstrap replicates.

### 2.1. PL25 Strain isolation, sequencing and assembly

Sediment samples were collected in 2023 from the hypersaline Pink Lake in Al Lith, Saudi Arabia (19.799183° N, 40.732525° E). At the sampling site, the water temperature was 33.5 °C, salinity was 220.7 PSU, conductivity was 274.7 mS/cm, pH was 6.68, dissolved oxygen saturation was 66.2% (1.44 mg/L), and chlorophyll a concentration was 0 µg/L. *V. salarius* PL25 was isolated and purified on marine medium at 37 °C. Genomic DNA was subsequently extracted from PL25 using a commercial DNA extraction kit, and its quantity and integrity were assessed using Qubit fluorometry and TapeStation electrophoresis. Long-read sequencing libraries were prepared using the Rapid Barcoding Kit V14 (SQK-RBK114-96; Oxford Nanopore Technologies, Oxford, UK) and sequenced on an Oxford Nanopore PromethION platform using an R10.4.1 flow cell (FLO-PRO114M). Raw signal data were base-called using Guppy v6.4.3 with the high-accuracy (HAC) model, and the resulting reads were assembled using Flye v2.9.3-b1797 (Kolmogorov et al., 2019) with the --nano-raw option. The circularity of contig_11 was assessed directly using the Nanopore reads. Briefly, the final 20 kb of the contig was joined to its first 20 kb to generate a reference sequence in which the assembler’s end-to-start junction was positioned at a defined location. The number of reads spanning this junction was then compared with the number spanning an equivalent position within an internal 40-kb control window. Additional details are provided in Supplementary Methods S1.

### 2.2. Bacterial strains and culture conditions

The antibacterial activity of PL25_M23 was evaluated against seven Gram-positive strains: *Streptococcus equi* subsp. *zooepidemicus* ATCC 43079, *Streptococcus pneumoniae* ATCC 49619, methicillin-resistant *Staphylococcus aureus* ATCC BAA-1026, methicillin-sensitive *S. aureus* subsp. *aureus* ATCC 29213, *Mammaliicoccus sciuri* ATCC 29061, *S. aureus* subsp. *aureus* ATCC BAA-976, and *V. salarius* PL25. *S. equi subsp. zooepidemicus* ATCC 43079, *S. pneumoniae* ATCC 49619, *S. aureus* ATCC BAA-1026, and *S. aureus* ATCC BAA-976 were cultured in brain heart infusion (BHI) broth and on BHI agar at 37 °C for 24 h. *S. pneumoniae* ATCC 49619 was incubated under 5% CO_2_, whereas the other three strains were incubated under aerobic conditions. *S. aureus* ATCC 29213 was cultured in tryptic soy broth (TSB) and on tryptic soy agar (TSA) at 37 °C for 24 h under aerobic conditions. *M. sciuri* ATCC 29061 was cultured in nutrient broth and on nutrient agar at 37 °C for 24 h under aerobic conditions. *V. salarius* PL25 was cultured in marine broth and on marine agar at 37 °C for 24 h under aerobic conditions.

### 2.3. Genome annotation

Whole-genome annotation was performed using Prokka v1.15.6 (Seemann, 2014), with gene prediction by Prodigal v2.6.3 (Hyatt et al., 2010). Prokka was run with the following parameters: --kingdom Bacteria --genus *Virgibacillus* --species *salarius* --strain PL25 --rfam --addgenes -- locustag PL25. The annotation identified 4,613 coding sequences (CDSs), 18 rRNAs, 66 tRNAs, and one tmRNA across eight contigs.

### 2.4. Plasmid identification

All eight contigs in the PL25 genome assembly were screened for plasmid-associated sequences using geNomad v1.7.0 (Camargo et al., 2024) in end-to-end mode against the geNomad v1.7 database. Relaxed marker-enrichment thresholds were applied (--min-virus-marker-enrichment 0 --min-plasmid-marker-enrichment 0 --min-virus-hallmarks 0 --min-plasmid-hallmarks 0) to improve the detection of divergent mobile genetic elements. Contig_11 (pPL25-1; 74,025 bp) was subsequently characterized using MOB-suite v3.1.9 (Robertson and Nash, 2018) with the mob_typer module. The completeness of its conjugation system was evaluated using MacSyFinder v2.1.6 (Néron et al., 2023) and the CONJScan v2.1.0 model set (Cury et al., 2020). Contig_11 was screened against all 17 plasmid conjugation models (Supplementary Methods S2). To identify a putative origin of transfer (oriT), motifs derived from IncP RP4 (Pansegrau and Lanka, 1991), IncQ RSF1010 (Becker and Meyer, 2000), IncW R388 (Llosa et al., 1994), pIP501 (Wang and Macrina, 1995), and ICEBs1 (Lee et al., 2010) were examined on both strands of contig_11. Specifically, inverted repeats of at least 18 bp were searched for within a ±3-kb region surrounding PL25_00076. Detailed motif information and command-line parameters are provided in Supplementary Methods S3, and the corresponding results are summarized in Supplementary Table S1.

To determine whether pPL25-1 could be assigned to an established plasmid taxonomic group, it was compared against two reference panels independent of the MOB-suite database used in the preceding analysis. First, k-mer-based Mash v2.3 sketch distances (Ondov et al., 2016) were calculated against the PLSDB 2025 dataset, which comprises 72,556 closed plasmids (Molano et al., 2025), using the released PLSDB Mash sketch with default settings. The six closest matches were subsequently examined by pairwise BLASTn analysis to quantify nucleotide-level sequence homology and identify the corresponding alignment coordinates on pPL25-1. Second, formal Plasmid Taxonomic Unit (PTU) classification was performed using COPLA v1.0 (Redondo-Salvo et al., 2021), with the bin/copla.py script and the Copla_RS84 reference graph, which comprises 9,894 plasmids. Classification was performed using the -t circular parameter and the full set of taxonomic priors: -k Bacteria, -p Bacillota, -c Bacilli, -o Bacillales, -f Bacillaceae, and -g *Virgibacillus*. COPLA’s auxiliary modules included MOBScan for relaxase typing (Garcillán-Barcia et al., 2020), PlasmidFinder for replicon typing (Carattoli et al., 2014), and the CARD database for antimicrobial resistance cargo screening (Alcock et al., 2023). PTU assignment required the query node to cluster within a Copla graph component containing at least four reference plasmids. Queries assigned to components containing fewer than four reference plasmids were classified as candidate novel PTUs.

### 2.5. Mobile elements and lysin domain identification

To identify mobile element marker genes on contig_11, the Prokka-predicted proteome was screened against a panel of Pfam and cell wall hydrolase hidden Markov models (HMMs), comprising 12 profiles, as described in Supplementary Methods S4. The analysis was performed using HMMER v3.4 (Eddy, 2011) and the hmmscan function, with an E-value threshold of ≤1 × 10^−5^. Pfam profiles were obtained from release 37.0 of the Pfam database, downloaded on May 14, 2026 (Mistry et al., 2021). A separate HMMER search was performed using a panel of Pfam profiles associated with lysin domains, including PF01551 (Peptidase_M23), PF00877 (NlpC/P60), PF00959 (GH24), PF01471 (PG_binding_1), PF01510 (Amidase_2), and PF01520 (Amidase_3). PL25_M23 (PL25_00078) and four additional mobile element-associated proteins (PL25_00076, PL25_00077, PL25_00082, and PL25_00148) were queried against the NCBI nonredundant (nr) protein database, accessed on May 17, 2026 (Sayers et al., 2022), using BLASTp from BLAST+ v2.16.0 (Camacho et al., 2009), with the parameters-evalue 1e-5 and - max_target_seqs 250. A circular map of pPL25-1 was generated from the Prokka GenBank annotation of contig_11 using pyCirclize v1.10.1 (https://github.com/moshi4/pyCirclize), together with matplotlib v3.10.9 (Hunter, 2007) and Biopython v1.83 (Cock et al., 2009).

### 2.6. Domain and signal peptide prediction

The domain architecture of PL25_M23 (PL25_00078) was annotated using InterProScan v5.61-93.0 (Blum et al., 2025) against multiple protein domain databases, including Pfam, the Conserved Domain Database (CDD), SUPERFAMILY, Gene3D, SMART, PROSITE, PRINTS, and PANTHER. Signal peptide prediction was performed using SignalP v6.0h (Teufel et al., 2022) with the --organism other parameter. In addition, a local Kyte–Doolittle hydropathy profile (Kyte and Doolittle, 1982), calculated using a sliding window of 19 residues, was generated in Python (Cock et al., 2009) as an independent assessment of potential non-cleavable N-terminal hydrophobic regions.

### 2.7. Phylogenetic analysis

To determine the phylogenetic position of PL25_M23 among characterized M23-family enzymes, we assembled a dataset comprising 101 M23-domain sequences. The dataset included PL25_M23, 80 high-scoring homologs retrieved from the NCBI nr database, five characterized reference M23 enzymes (lysostaphin, UniProt P10547; LytM, O33599; ALE-1, O05156; EnpA, E1V3I0; and zoocin A, O54309; UniProt accessed on May 18, 2026) (The UniProt Consortium et al., 2023), the 314-amino-acid catalytic core of CwlP (residues 1373–1686 of UniProt O31976), and 14 phage-associated M23 lysins. CwlT (UniProt P96645) was excluded because its C-terminal peptidase domain belongs to the NlpC/P60 cysteine peptidase family (Pfam PF00877). Sequences were aligned using MAFFT v7.526 (--auto) (Katoh and Standley, 2013), and poorly aligned columns were removed using trimAl v1.5.rev1 (-gappyout) (Capella-Gutiérrez et al., 2009). Maximum-likelihood phylogenetic reconstruction was performed using IQ-TREE v3.1.1 with the parameters -m MFP -B 1000 -alrt 1000 (Minh et al., 2020). ModelFinder selected the Q.PFAM+G4 substitution model using the Bayesian information criterion (Kalyaanamoorthy et al., 2017; Minh et al., 2021). Branch support was assessed using 1,000 ultrafast bootstrap replicates (Hoang et al., 2018) and 1,000 SH-aLRT replicates (Anisimova et al., 2011). The resulting tree was midpoint-rooted and visualized using Bio.Phylo (Cock et al., 2009).

To identify PL25_M23 homologs encoded in the closed genomes of salt-adapted taxa, the 250 BLASTp hits retained from the initial search were first screened against a predefined list of 15 halophilic or halotolerant genera: *Halobacillus*, *Lentibacillus*, *Virgibacillus*, *Gracilibacillus*, *Oceanobacillus*, *Cytobacillus*, *Ornithinibacillus*, *Salinicoccus*, *Salimicrobium*, *Salibacillus*, *Salinibacillus*, *Pontibacillus*, *Aquibacillus*, *Halolactibacillus*, and *Salirhabdus*. This taxonomic filter reduced the dataset to 44 hits. Protein records were linked to their corresponding genome assemblies through NCBI Identical Protein Groups (IPG), and only RefSeq assemblies with GCF_ accessions and an AssemblyStatus of “Complete Genome” or “Chromosome” were retained. The organism associated with each assembly was also required to belong to the predefined genus list, and only one protein sequence was retained per assembly. No additional filtering was applied based on sequence identity, alignment coverage, protein length, or isolation environment. This procedure yielded 14 homologs from closed RefSeq assemblies.

Replicon type was assigned based on the title of the corresponding RefSeq nucleotide record linked through the IPG entry. Records with titles containing “plasmid” were classified as plasmid-encoded, whereas those containing “chromosome” or “complete genome” were classified as chromosomal. Host and strain designations were standardized based on the corresponding nucleotide-record titles, and plasmid names were manually curated when automated parsing was ambiguous. For the focused phylogenetic analysis, full-length protein sequences of the 14 homologs, PL25_M23, and lysostaphin (UniProt P10547) were aligned using MAFFT v7.526 with the parameters --auto --thread 1. The resulting alignment was trimmed using trimAl v1.5.rev1 (-gappyout). Maximum-likelihood phylogenetic reconstruction was performed using IQ-TREE v3.1.1 with ModelFinder (-m MFP) (Kalyaanamoorthy et al., 2017; Minh et al., 2020). ModelFinder selected WAG+G4 according to the Bayesian information criterion (Whelan and Goldman, 2001). Branch support was assessed using 1,000 ultrafast bootstrap replicates and 1,000 SH-aLRT replicates. The resulting tree was rooted using lysostaphin as the outgroup and visualized using Bio.Phylo.

### 2.8. Structure prediction and comparison

To compare the predicted structure of PL25_M23 with those of characterized homologs, structural models of PL25_M23 (residues 38–370) and the SLT/M23 catalytic core of CwlP (residues 1373– 1686; UniProt O31976) were generated using ColabFold v1.5.5 (Mirdita et al., 2022) with the AlphaFold2-ptm model, five models, and three recycles (Jumper et al., 2021). Reference structures were retrieved from the RCSB Protein Data Bank (lysostaphin, 4QPB; LytM, 4ZYB; and zoocin A, 5KVP; PDB accessed on May 20, 2026) and the AlphaFold Protein Structure Database (ALE-1, AF-O05156-F1 v6; and CwlT, AF-P96645-F1 v4; accessed on May 22, 2026) (Varadi et al., 2022). Pairwise structural superpositions were performed using PyMOL v3.1.6.1 (Open-Source) (Schrödinger, 2024) with the cmd.super() function, using five cycles and a 2.0-Å outlier rejection cutoff. Domain-restricted superpositions were performed according to the domain boundaries defined by InterProScan. The corresponding root-mean-square deviations (RMSDs) and numbers of aligned residues are reported in Supplementary Table S2.

### 2.9. Sequence charge and surface comparison

To compare the surface charge characteristics of PL25_M23 and reference hydrolases, the theoretical isoelectric point (pI) and net charge at pH 7.0 were calculated for each protein using the Protein Analysis module of Biopython v1.83 (Cock et al., 2009) based on the Henderson– Hasselbalch model. The sequences used for these calculations and structural modeling comprised PL25_M23 residues 38–370 (333 amino acids), the CwlP catalytic core corresponding to residues 1373–1686 of UniProt O31976 (314 amino acids), the full-length CwlT sequence represented by the AlphaFold2 model of UniProt P96645 (329 amino acids), lysostaphin (chain A of PDB entry 4QPB; 133 amino acids), and LytM (chain A of PDB entry 4ZYB; 133 amino acids). Net charge density (%) was calculated using the following equation: (K + R − D − E)/(total number of residues) × 100, where K, R, D, and E represent lysine, arginine, aspartic acid, and glutamic acid residues, respectively. Surface representations were generated using PyMOL v3.1.6.1 (Open-Source) (Schrödinger, 2024), with a consistent per-atom side-chain coloring scheme applied to all five proteins. The corresponding parameters are provided in Supplementary Methods S5.

### 2.10. Recombinant protein expression and purification

For functional assays, a synthetic construct encoding PL25_M23 residues 44–370, preceded by an N-terminal Met-His_6_ tag (MHHHHHH; without a linker or protease cleavage site), was generated. The resulting recombinant protein comprised 334 amino acids, with a calculated molecular mass of 36.5 kDa and a theoretical pI of 4.83. The construct was cloned into the NdeI and HindIII sites of pET-30a, and recombinant protein production was performed under contract by GenScript (Piscataway, NJ, USA; order U054ZZPXG0, lot U054ZZPXG0-4/P18LA001). The construct begins at residue 44, thereby omitting most of the predicted N-terminal hydrophobic region spanning residues 19–46. SignalP v6.0h did not predict a cleavable signal peptide. The recombinant protein was expressed in *Escherichia coli* BL21 Star™ (DE3), recovered from inclusion bodies, and purified using nickel-nitrilotriacetic acid (Ni-NTA) affinity chromatography. The purification yielded 41.76 mg of protein distributed across four 4.0-mL aliquots. Protein refolding was performed by direct dialysis from a denaturing buffer containing 50 mM Tris-HCl and 8 M urea (pH 8.0) into the final storage buffer containing 50 mM Tris-HCl, 150 mM NaCl, and 10% glycerol (pH 8.0). The purified protein was supplied in the non-denaturing storage buffer (50 mM Tris-HCl, 150 mM NaCl, 10% glycerol, pH 8.0) at a concentration of 2.61 mg/mL, as determined using the Bradford assay (Bradford, 1976). The protein was ≥90% pure, as assessed by SDS-PAGE under reducing conditions, and produced a prominent band of the expected molecular mass in an anti-His immunoblot (Supplementary Figure 1). No visible precipitation was observed following one freeze–thaw cycle. Protein aliquots were stored at −80 °C and thawed once before use.

### 2.11. Antibacterial activity assays

#### Bacterial strains and preparation

Bacterial strains were prepared under the strain-specific culture conditions described above. Cultures were harvested, washed with assay buffer, and adjusted to approximately 5 × 10^8^ CFU/mL before being added to the assay mixtures, yielding a final reaction inoculum of approximately 2.5 × 10^8^ CFU/mL after 1:1 mixing. All PL25_M23 activity assays were conducted in a total reaction volume of 200 µL in PBS (pH 7.4) at 37 °C. PL25_M23 was added from the purified protein stock to achieve the indicated final concentration. Untreated controls contained the corresponding bacterial suspension without PL25_M23. In addition to the untreated control, a storage-buffer control was included in which bacteria received the same volume of protein storage buffer (50 mM Tris-HCl, 150 mM NaCl, 10% glycerol, pH 8.0) as the corresponding PL25_M23-treated reaction, but without the enzyme. Because PL25_M23 was added from a single stock at 2.61 mg/mL, the volume of storage buffer carried over into the reaction increased with the nominal enzyme concentration, accounting for 1.9%, 3.8%, and 11.5% (v/v) of the 200-µL reaction volume at 50, 100, and 300 µg/mL, respectively. Storage-buffer controls were included in the primary antibacterial assay at 100 µg/mL and across the full dose-response series. Unless otherwise stated, all PL25_M23, EDTA, ZnCl_2_, and additional NaCl concentrations refer to their final concentrations in the 200-µL reaction. Viable counts were determined using the Miles–Misra drop-plate method (Miles et al., 1938). Reactions were serially diluted 10-fold in PBS, and 10-µL drops from appropriate dilutions were plated onto the corresponding agar medium and incubated under the strain-specific conditions described above. Colony counts within the countable range were used to calculate CFU/mL as follows: colony count × dilution factor × 100.

#### Termination of enzymatic activity by dilution

At each sampling point, the reaction was removed from 37 °C and immediately serially diluted 10-fold in room-temperature PBS. This procedure reduced the PL25_M23 concentration by at least 10-fold in the first dilution step, with further reductions at each subsequent step, before the drops were plated and allowed to dry. The adequacy of this procedure was evaluated using the 0-min time point of the time-kill assay, in which bacteria were mixed with PL25_M23 at 100 µg/mL and immediately diluted and plated without incubation. Across the seven strains, this condition yielded a mean Δlog_10_ CFU/mL of −0.05 (95% CI, −0.14 to +0.04; P = 0.23; n = 21), which was not significantly different from the matched untreated control. These results indicate that additional killing during dilution, plating, and drying did not measurably contribute to the reductions reported here.

#### Dose response and selection of the working concentration

PL25_M23 was tested at concentrations of 0, 50, 100, and 300 µg/mL for 60 min at 37 °C. A concentration of 100 µg/mL was selected as the working concentration for subsequent kinetic and biochemical assays because it produced a mean reduction of 2.67 log_10_ CFU/mL across the strain panel while remaining below the highest concentration tested.

#### Antibacterial assay

PL25_M23 was tested at the selected working concentration of 100 µg/mL for 60 min at 37 °C against each strain. Matched PBS-only controls were run in parallel to establish the endpoint reductions in log_10_ CFU/mL.

#### Time-kill kinetics

PL25_M23 at 100 µg/mL was mixed with the bacterial suspensions as described above. Reactions were sampled at 0, 30, 60, 120, and 180 min and immediately diluted for viable-count determination using the Miles–Misra method. Control samples contained bacteria mixed 1:1 with PBS and were sampled at the same time points.

#### Metal-dependence assay

PL25_M23 was tested at 100 µg/mL in the presence or absence of 1 mM EDTA and/or 1 mM ZnCl_2_. The experimental conditions comprised untreated bacteria, 1 mM EDTA alone, 1 mM ZnCl_2_ alone, PL25_M23 alone, PL25_M23 + EDTA, PL25_M23 + ZnCl_2_, and PL25_M23 + EDTA + ZnCl_2_. Reactions were incubated for 60 min at 37 °C before viable counts were determined. EDTA-mediated inhibition was assessed by comparing PL25_M23 alone with PL25_M23 + EDTA, whereas restoration by Zn^2+^ was assessed by comparing PL25_M23 + EDTA with PL25_M23 + EDTA + ZnCl_2_.

#### Salt-response assay

PL25_M23 activity was assessed at 100 µg/mL in standard PBS supplemented with 0, 0.25, 0.5, or 1.0 M additional NaCl. Preliminary salt-only viability controls revealed pronounced losses in viability for *S. equi* subsp. *zooepidemicus* ATCC 43079 and *S. pneumoniae* ATCC 49619. At 1.0 M added NaCl, untreated viability decreased by approximately 4.8 and 4.1 log_10_ CFU/mL, respectively. These two strains were therefore excluded from the enzyme salt-response analysis to avoid confounding salt-induced losses in viability with PL25_M23 activity. The remaining five strains were evaluated across the full NaCl concentration range. Untreated controls in PBS supplemented with 0 or 1.0 M additional NaCl were included to verify viability at the two extremes of the tested range. Among the five retained strains, untreated viability did not differ significantly between 0 and 1.0 M added NaCl (mean difference, −0.05 log_10_ CFU/mL; not significant for any strain after Holm correction). Accordingly, the untreated control at 0 M added NaCl was used as the reference for all four salt levels, and salt-response data were expressed as Δlog_10_ CFU/mL. Because untreated controls were included only at 0 and 1.0 M added NaCl, the values at 0.25 and 0.5 M were referenced to the 0-M control without corresponding salt-matched untreated controls. The underlying viability data and the controls included in the experiment are presented in Supplementary Figure 2e.

#### Replication and statistical analysis

All experiments were performed using three independent biological replicates. Each biological replicate represented an independent experiment conducted on a separate day using a bacterial culture grown from a separate colony and a separate dilution of the enzyme stock. Within each biological replicate, each condition was prepared as two separate technical reaction replicates. The two technical CFU/mL measurements were averaged to obtain one value per condition for each biological replicate; these technical replicates were not treated as independent observations in the statistical analyses. CFU/mL values were log_10_-transformed before analysis. For matched antibacterial comparisons, killing was expressed as Δlog_10_ CFU/mL = log_10_(CFU/mL untreated) − log_10_(CFU/mL treated), with the difference calculated separately for each biological replicate. Data are presented as individual biological replicates, with mean ± SD where appropriate. Storage-buffer controls were analyzed in the same manner. Within each biological replicate, the buffer-control count was compared with its matched untreated count, and the resulting Δlog_10_ values were tested against zero at each buffer volume using one-sample t-tests, with Holm correction across the four volumes (*n* = 21 per volume; seven strains × three biological replicates).

#### Primary antibacterial assay at 100 µg/mL

Treated and matched untreated values were compared within each strain using paired t-tests. Dose-response and time-kill datasets were analyzed as Δlog_10_ CFU/mL reductions using repeated-measures fixed-effect models. Strain, concentration or time, and their interaction were included as fixed effects, with biological replicate included as a fixed blocking factor within strain. Contrasts between each concentration or time point and the corresponding zero level were estimated within each strain using *emmeans* (Lenth and Piaskowski, 2017). For the metal-dependence assay, the prespecified contrasts were PL25_M23 versus PL25_M23 + EDTA and PL25_M23 + EDTA versus PL25_M23 + EDTA + ZnCl_2_. These comparisons were assessed using paired t-tests.

#### Salt-response assay

Within each of the five retained strains, PL25_M23-treated counts at 0.25, 0.5, and 1.0 M added NaCl were compared with the corresponding PL25_M23-treated condition at 0 M additional NaCl using paired t-tests, yielding 15 planned comparisons. P-values were adjusted across these comparisons using the Holm method. These comparisons tested whether killing changed with increasing NaCl concentrations. Because the same 0-M untreated control was subtracted from both members of each contrast, the comparisons were equivalent to comparing the residual PL25_M23-treated counts across salt levels. At 1.0 M added NaCl, for which an untreated control was also available, a salt-matched reduction was additionally calculated within each biological replicate and compared with the salt-matched reduction at 0 M using paired t-tests, with Holm correction across the five strains. P-values were adjusted using the Holm method (Holm, Sture, 1979) across the planned comparisons within each figure panel. Statistical significance was defined as follows: ****P ≤ 0.0001, ***P ≤ 0.001, **P ≤ 0.01, *P ≤ 0.05, and ns, P > 0.05. Analyses were performed in R 4.5.1 (R Core Team, 2025) using *emmeans* 2.0.3 (Lenth and Piaskowski, 2017), together with the indicated data-processing and visualization packages.

## 3. Results

### 3.1. A 74-kb circular plasmid candidate in *V. salarius* PL25 encodes PL25_M23

The 4.75-Mb genome of *V. salarius* PL25 was annotated using Prokka. Genome-based taxonomic analysis assigned the isolate to *V. salarius*, with a dDDH value of 95.8% relative to the type strain of *V. salarius*, substantially exceeding the 70% species delineation threshold. The genome assembly comprised eight contigs (Supplementary Table S3), of which contig_4 (4,349,560 bp; 36.81% GC) was identified as the chromosome and contig_11 (74,025 bp) as a plasmid candidate, designated pPL25-1. Flye identified pPL25-1 as a closed circular sequence, with a mean coverage of 889× and an estimated copy number of two, compared with 476× coverage for contig_4. Circularity was further assessed directly using the Nanopore reads. A total of 608 reads aligned across the assembler-defined end-to-start junction, with at least 1 kb of aligned sequence on either side, compared with 516 reads spanning an equivalent internal position. The corresponding sequencing depths were 909× and 828×, respectively. This difference was maintained when more stringent alignment-length thresholds were applied, with 443 versus 392 reads at 2 kb and 218 versus 200 reads at 5 kb. Individual reads measuring 13–25 kb also aligned continuously across the junction. Consistent with these findings, geNomad assigned pPL25-1 a plasmid classification score of 0.989, supporting its classification as a circular plasmid candidate (Figure 1a; Supplementary Table S3). To identify mobile-element and cell wall hydrolase genes on pPL25-1, the predicted proteome was screened using HMMER against a panel of 12 Pfam profiles (Supplementary Methods S4). This analysis identified PL25_00078 as a 370-amino-acid Peptidase_M23 protein (PF01551; E = 1.5 × 10^−36^), designated PL25_M23. Three additional mobile-element-associated markers were identified: PL25_00082, encoding a VirB4-like AAA-domain ATPase (AAA_10; E = 8.4 × 10^−69^); PL25_00094, encoding a RecD2-like helicase (AAA_19; E = 1.7 × 10^−41^); and PL25_00148, encoding a T4SS-DNA-transfer-family protein (PF02534; E = 2.1 × 10^−12^). BLASTp analysis further supported the annotation of PL25_00076 as a replication–relaxation-family protein (top hit: *Ornithinibacillus contaminans* WP_157052516.1; 57% identity; E = 3.6 × 10^−135^) and PL25_00077 as a RusA-family crossover-junction endodeoxyribonuclease (top hit from the same *O. contaminans* genome; 58% identity; E = 6.0 × 10^−58^). These five putative transfer- or maintenance-associated proteins were located in the same region of pPL25-1 as PL25_M23 (Figure 1a). Compositional analysis further distinguished pPL25-1 from the chromosome. The overall GC content of pPL25-1 was 31.22%, compared with 36.81% for contig_4 (Δ = −5.59%). A similar difference was observed at the third codon position: GC3 was 23.19% for pPL25-1 and 28.71% for contig_4 (Δ = −5.52%) (Figure 1b). MOB-suite classified pPL25-1 as non-mobilizable and detected neither a recognized MOB-family relaxase nor a complete mating-pair formation system (Supplementary Table S1) (Robertson and Nash, 2018).

**Figure 1.**
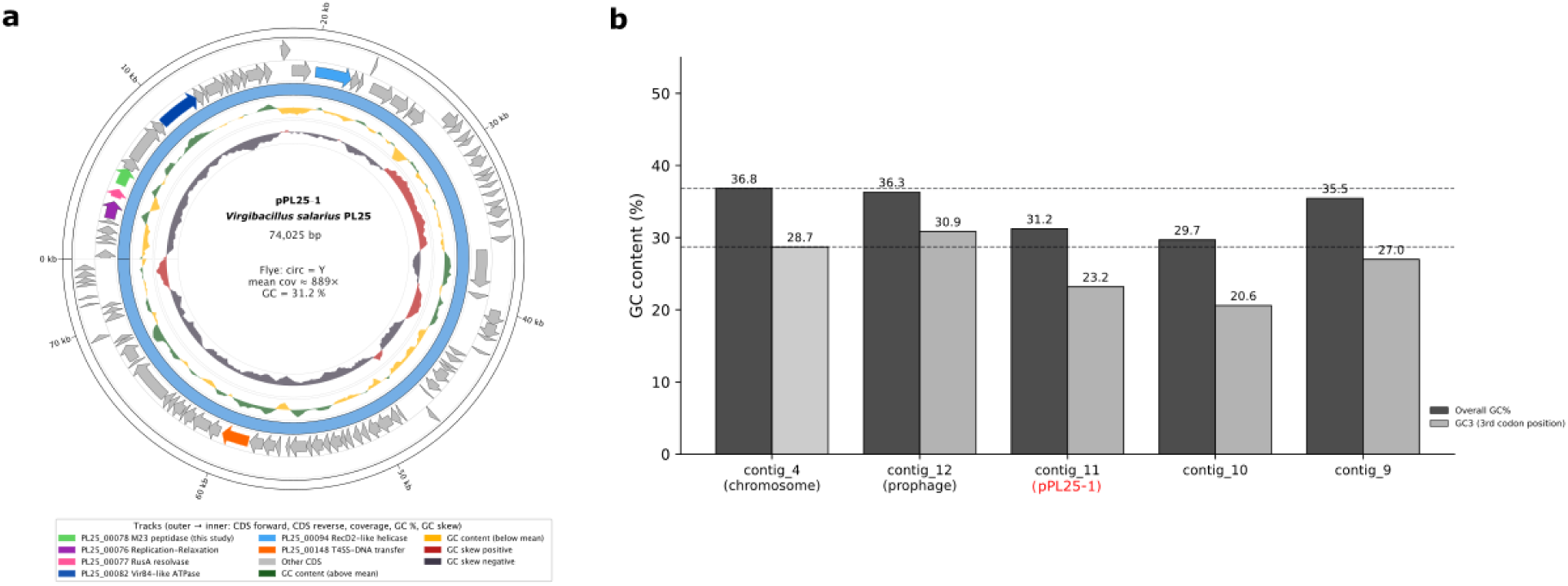
Circular map of pPL25-1 carrying PL25_M23 and its GC content. (a) Circular map of pPL25-1 (contig_11; 74,025 bp) generated using pyCirclize based on the Prokka annotation. From the outermost to the innermost track, the map shows forward-strand CDS arrows, reverse-strand CDS arrows, per-base sequencing coverage, GC-content deviation, and GC skew. PL25_M23 and five putative mobile-element-associated proteins are highlighted: PL25_00076 (replication–relaxation-family protein, purple), PL25_00077 (RusA-family resolvase, pink), PL25_00078 (PL25_M23, the predicted dual-domain SLT/M23 hydrolase, green), PL25_00082 (VirB4-like ATPase, dark blue), PL25_00094 (RecD2-like helicase, light blue), and PL25_00148 (T4SS-DNA-transfer-family protein, orange). All other CDSs are shown in gray. The plot origin was set at the start of PL25_00076, following the standard convention for circular plasmid maps. (b) Overall GC content (dark bars) and GC content at the third codon position (GC3; light bars) for the five annotated PL25 contigs. pPL25-1 has lower overall GC content and GC3 than the main chromosome, indicating a distinct compositional profile. Three unannotated contigs were excluded from this panel: contig_2 (a 249-kb chromosomal repeat), contig_5 (462 bp), and contig_6 (514 bp).

### 3.2. PL25_M23 is associated with plasmid-encoded homologs in closed genomes

To place PL25_M23 within the M23 family and determine whether related sequences occur in plasmid or chromosomal contexts, we searched the NCBI nr protein database using BLASTp. All 250 hits retained for analysis were annotated as M23-family peptidoglycan DD-metalloendopeptidases. The two closest homologs were WP_047980627.1 from *Ornithinibacillus contaminans* (66.6% amino acid identity; E = 2.1 × 10^−158^) and WP_066194568.1 from *Cytobacillus horneckiae* (64.0% identity; E = 6.6 × 10^−153^). Most of the remaining hits originated from unresolved whole-genome shotgun (WGS) contigs. The maximum-likelihood phylogeny comprising 101 M23-family sequences placed PL25_M23 within the Bacillaceae subclade, most closely associated with WP_047980627.1 (*O. contaminans*) and WP_066194568.1 (*C. horneckiae*). PL25_M23 was separated from the characterized reference M23 enzymes (lysostaphin, LytM, ALE-1, EnpA, zoocin A, and CwlP) and the 14 phage-associated M23 lysins (Figure 2; 101-taxon topology in Supplementary Figure 3).

**Figure 2.**
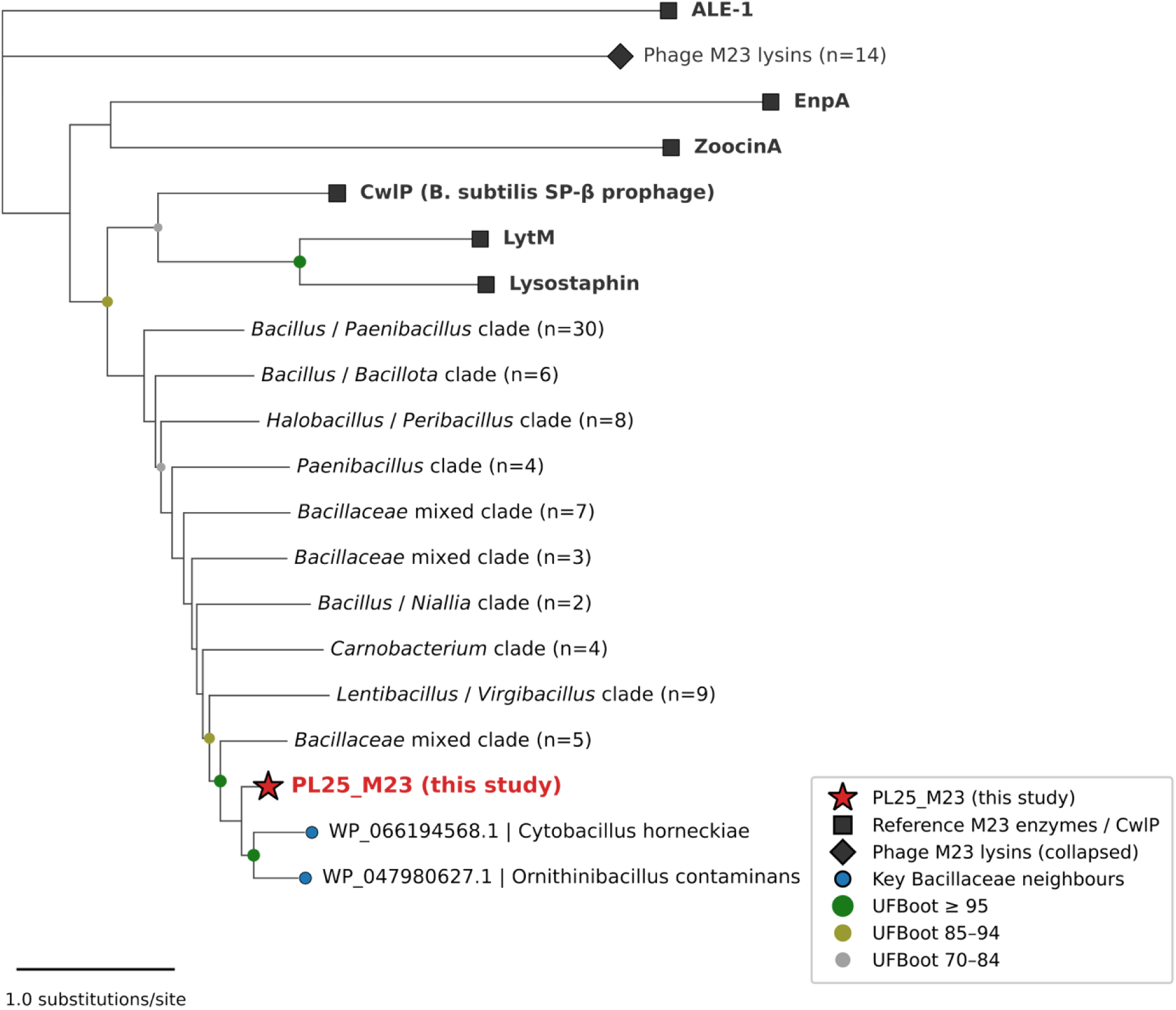
Maximum-likelihood phylogeny of PL25_M23 and related M23-family hydrolases. The tree includes 101 M23-family sequences: PL25_M23, 80 NCBI nr homologs, five reference M23 enzymes, 14 phage-associated M23 lysins, and the catalytic core of CwlP. The tree was inferred using IQ-TREE v3.1.1 under the Q.PFAM+G4 model, with 1,000 ultrafast bootstrap and 1,000 SH-aLRT replicates, and was midpoint-rooted. PL25_M23 (red star) is positioned adjacent to homologs from *O. contaminans* and *C. horneckiae* and is separated from the reference M23 enzymes (purple squares), CwlP (purple star), and phage-associated M23 lysins (orange diamonds). Node labels indicate SH-aLRT/UFBoot support values. Branch and taxon-label colors indicate the categories defined in the figure.

Among the 14 PL25_M23 homologs retained from the predefined set of halophilic or halotolerant genera, which spanned six genera, 11 were encoded on plasmids and three were chromosomally encoded (Supplementary Table S4; Supplementary Figure 4). These homologs were selected based on genus membership, RefSeq status, and closed-assembly status, without applying a sequence-identity threshold. Their amino acid identities to PL25_M23 ranged from 35.0% to 53.0%. The two highest-identity hits in the full BLASTp search, *O. contaminans* WP_047980627.1 (66.6%) and *C. horneckiae* WP_066194568.1 (64.0%), were not included in this focused dataset because their source assemblies did not meet the closed-assembly criterion. A focused maximum-likelihood phylogeny comprising the 14 retained homologs, PL25_M23, and lysostaphin recovered a well-supported plasmid-associated clade containing nine homologs (SH-aLRT = 98.8; UFBoot = 99) and a distinct chromosomal clade comprising sequences from *Lentibacillus* and *V. ainsalahensis* (SH-aLRT = 99.9; UFBoot = 99). PL25_M23 was positioned adjacent to the plasmid-associated clade. The nine homologs in this clade ranged from 363 to 758 amino acids in length, whereas PL25_M23 (370 amino acids) and the remaining five homologs (358–426 amino acids) were substantially shorter. This length distribution suggests that PL25_M23 belongs to a short-form subgroup of Bacillaceae M23 hydrolases that occurs in both plasmid and chromosomal contexts.

Mash sketch-distance analysis against the PLSDB 2025 dataset (*n* = 72,556 closed plasmids) identified no neighbors with a distance of d ≤ 0.2. The closest hit was the *Virgibacillus* sp. Bac332 plasmid NZ_CP033047.1 (d = 0.263; 2/1,000 shared k-mers). Seventeen additional hits had a distance of d = 0.296 (1/1,000 shared k-mers), whereas the remaining 72,538 plasmids shared no k-mers with pPL25-1. Pairwise BLASTn analysis of the six closest hits yielded alignments only for NZ_CP033047.1. These alignments comprised 302 bp across three blocks (0.40% of the pPL25-1 sequence; 97.4% identity), all located within a single hypothetical CDS (PL25_00127), with no overlap with PL25_M23 or the five putative mobile-element-associated genes. COPLA v1.0 classification against the Copla_RS84 graph (*n* = 9,894 plasmids) placed pPL25-1 in a graph component containing only one sequence, below the minimum component size of four required for PTU assignment. Accordingly, COPLA classified pPL25-1 as a candidate “new, still unnamed, PTU” (score = 1.0). COPLA’s auxiliary modules detected no MOBscan relaxase, PlasmidFinder replicon, or CARD-annotated antimicrobial resistance or virulence cargo. Similarly, MacSyFinder/CONJScan did not identify any complete conjugation systems (Supplementary Table S1). Collectively, these findings indicate that pPL25-1 has no close relative among the closed plasmid sequences represented in the databases examined.

### 3.3. PL25_M23 has a predicted dual-domain SLT/M23 architecture

To characterize the domain composition of PL25_M23, the protein sequence was analyzed using five domain-annotation databases. Two putative catalytic domains were identified within the 370-amino-acid sequence. The N-terminal region was assigned an Slt35-like/lysozyme glycan-hydrolase fold by CDD cd13399 (residues 86–214), SUPERFAMILY SSF53955 (residues 70– 222), and Gene3D G3DSA:1.10.530.10 (residues 68–229). The C-terminal region was identified as a Peptidase_M23 domain by Pfam PF01551 (residues 263–360; E = 2.3 × 10^−32^), CDD cd12797 (residues 264–351), and PANTHER PTHR21666 (residues 203–365). SignalP classified the sequence as “OTHER” and did not predict a cleavable signal peptide. Kyte–Doolittle hydropathy analysis identified a hydrophobic stretch spanning residues 19–46 (Figure 3a). The AlphaFold2 model of PL25_M23 residues 38–370, with a mean pLDDT of 88.7, predicted a two-domain architecture consisting of distinct SLT- and M23-domain lobes connected by a linker region (Figure 3b). The M23 peptidase domain contains the HxxxD motif at residues H264–D268 and the HxH motif at residues H346–H348 (Figure 3b; Supplementary Table S5).

**Figure 3.**
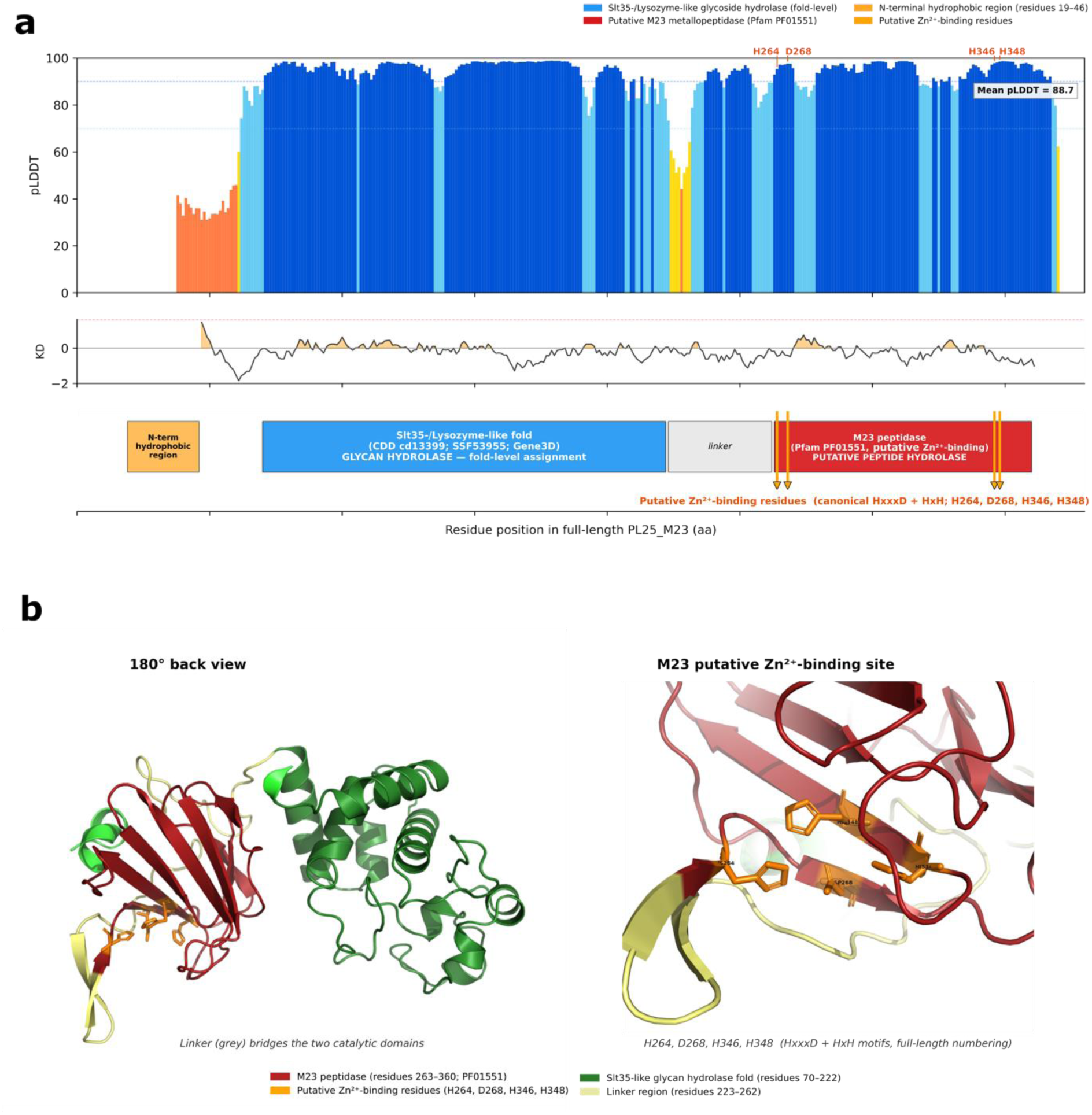
Predicted dual-domain architecture of PL25_M23. (a) Domain architecture and AlphaFold2 structure of PL25_M23 residues 38–370. InterProScan identified an N-terminal Slt35-/lysozyme-like fold (residues 68–229) and a C-terminal Peptidase_M23 domain (PF01551; residues 263–360). The Peptidase_M23 domain contains the putative Zn^2+^-binding motifs HxxxD and HxH at residues H264–D268 and H346–H348, respectively. The Kyte–Doolittle hydropathy profile, displayed below the pLDDT bars, indicates a potentially hydrophobic N-terminal region spanning residues 19–46, which is highlighted in orange. (b) AlphaFold2 structure of PL25_M23 residues 38–370, with a mean pLDDT of 88.7. Views rotated by 90° and 180° show the predicted two-domain architecture. The close-up highlights H264, D268, H346, and H348, with residue positions given according to the full-length sequence.

### 3.4. Structural superposition supports similarity between the PL25_M23 and CwlP catalytic domains

To assess whether the predicted catalytic domains of PL25_M23 adopt folds similar to those of CwlP, we superposed the AlphaFold2 model of PL25_M23 onto the CwlP catalytic core and CwlT, restricting the structural alignments to individual domains. For the C-terminal peptidase domain (Figure 4a), PL25_M23 aligned with the M23 domain of CwlP with an RMSD of 0.69 Å over 117 aligned residues. In contrast, its alignment with the NlpC/P60 peptidase domain of CwlT yielded an RMSD of 3.23 Å over 16 aligned residues. For the N-terminal glycan-hydrolase domain (Figure 4b), PL25_M23 aligned with the SLT domain of CwlP with an RMSD of 3.31 Å over 108 aligned residues, whereas alignment with the GH23-like domain of CwlT yielded an RMSD of 10.43 Å over 109 aligned residues. When the full-length two-domain regions were superposed without restricting the alignments to individual domain boundaries (Figure 4c), PL25_M23 aligned with CwlP with an RMSD of 16.75 Å over 234 aligned residues and with CwlT with an RMSD of 10.43 Å over 109 aligned residues. Thus, the domain-restricted alignments support structural similarity between the corresponding catalytic domains of PL25_M23 and CwlP. The substantially higher RMSD obtained from the unrestricted alignment suggests that the relative domain arrangement differs between the proteins, that the predicted interdomain orientation is uncertain, or both (Jumper et al., 2021).

**Figure 4.**
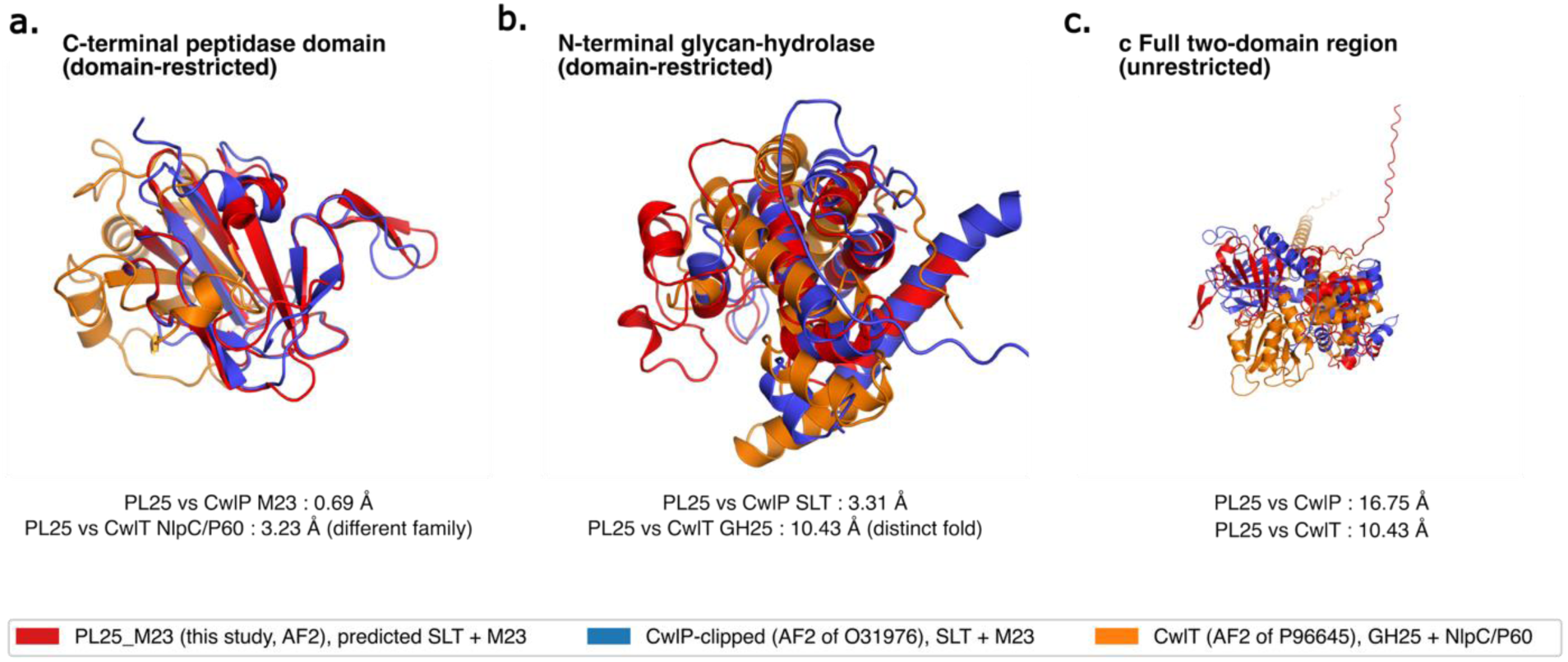
Structural comparison of PL25_M23, CwlP, and CwlT. (a) Structural alignment restricted to the C-terminal peptidase domain. The PL25_M23 M23 domain aligned with the CwlP M23 domain with an RMSD of 0.69 Å across 117 aligned residues. CwlT is included for architectural comparison only. PL25_M23 is shown in red, the truncated catalytic core of CwlP in blue, and CwlT in orange. (b) Domain-restricted structural alignment of the N-terminal glycan-hydrolase regions. The Slt35-/lysozyme-like fold of PL25_M23 aligned with the CwlP SLT domain, with an RMSD of 3.31 Å across 108 aligned residues. CwlT has a distinct GH23-like fold and is included for architectural comparison only. (c) Unrestricted structural alignment of the predicted two-domain regions, which yielded an RMSD of 16.75 Å across 234 aligned residues.

### 3.5. PL25_M23 has an acidic predicted surface

To determine whether PL25_M23 exhibits the electrostatic characteristics commonly reported for halophilic proteins, we compared its predicted surface charge profile with those of four reference hydrolases. Sequence-based analysis of the modeled PL25_M23 region (residues 38–370) yielded a pI of 4.54 and a net charge of −22.4 at pH 7. The acidic and basic residue contents were 12.3% and 5.4%, respectively, resulting in a net charge density of −6.9% (Figure 5b). For the four comparator proteins, the corresponding pI, net charge, and net charge density values were as follows: CwlP catalytic core, 9.76, +19.1, and +6.1%; full-length CwlT, 5.11, −12.9, and −3.6%; lysostaphin (PDB 4QPB), 9.11, +3.4, and +2.3%; and LytM (PDB 4ZYB), 8.18, +1.3, and +0.8% (Figure 5b). Per-atom surface representations showed the distribution of acidic side chains across the front and 180°-rotated back views of PL25_M23 (Figure 5a). In contrast, the comparator proteins displayed a mixture of acidic and basic surface patches. Among the proteins examined, PL25_M23 therefore had the most acidic predicted surface-residue profile. This compositional feature is consistent with patterns reported for halophilic proteins, although it is not specific to halophilic adaptation and may also arise under other selective pressures.

**Figure 5.**
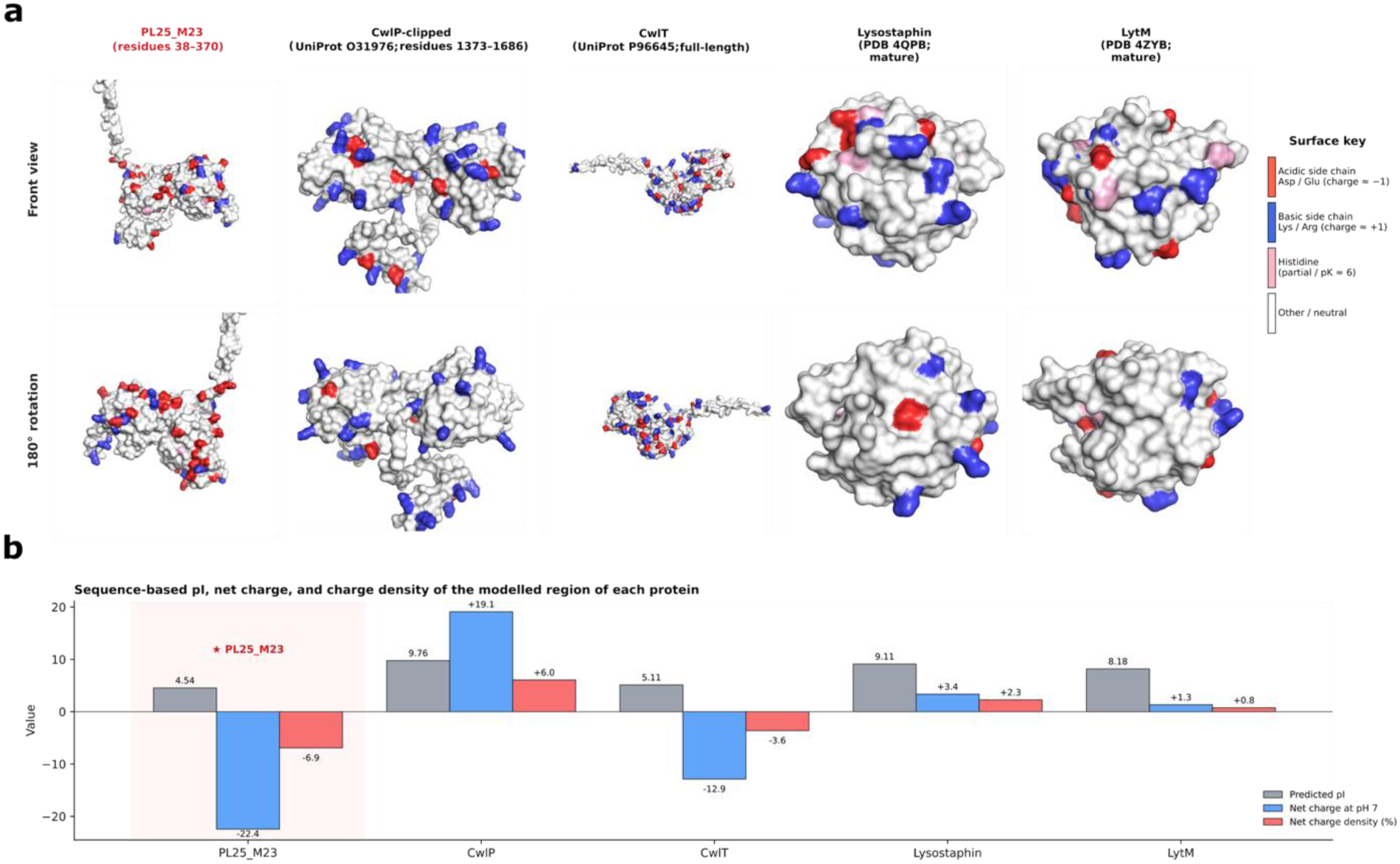
Predicted acidic surface profile of PL25_M23. (a) Surface distribution of charged side chains in PL25_M23 and four reference proteins (CwlP, CwlT, lysostaphin, and LytM), shown in front and 180°-rotated views. Acidic side chains (Asp and Glu) are shown in red, basic side chains (Lys and Arg) in blue, and histidine residues in pink. PL25_M23 and the CwlP and CwlT comparators are represented by AlphaFold2 models, whereas lysostaphin (PDB 4QPB) and LytM (PDB 4ZYB) are represented by experimentally determined structures. (b) Sequence-based pI, net charge at pH 7.0, and net charge density for the modeled or experimentally resolved region of each protein. Among the protein s compared, PL25_M23 had the lowest predicted pI (4.54), the most negative predicted net charge at pH 7.0 (−22.4), and the most negative charge density (−6.9%).

### 3.6. PL25_M23 reduces viable counts across a seven-strain Gram-positive panel

Recombinant PL25_M23 exhibited antibacterial activity against all seven Gram-positive strains tested. At a concentration of 100 µg/mL and an exposure time of 60 min, the mean reductions in viable counts ranged from approximately 1.93 log_10_ CFU/mL for *S. pneumoniae* ATCC 49619 to 3.78 log_10_ CFU/mL for *V. salarius* PL25. Intermediate reductions were observed for *S. equi* subsp. *zooepidemicus* ATCC 43079, MRSA *S. aureus* ATCC BAA-1026, MSSA *S. aureus* ATCC 29213, *M. sciuri* ATCC 29061, and *S. aureus* ATCC BAA-976 (Figure 6a). Antibacterial activity increased with PL25_M23 concentration across the 0–300 µg/mL range, with mean viable-count reductions of 0.82 log_10_ CFU/mL at 50 µg/mL, 2.67 log_10_ CFU/mL at 100 µg/mL, and 3.25 log_10_ CFU/mL at 300 µg/mL (Figure 6b). Time-kill analysis at 100 µg/mL showed that most of the reduction in viable counts occurred within the first 30–60 min, followed by smaller additional reductions through 120–180 min in several strains (Figure 6c). The largest reductions were observed for *V. salarius* PL25 and MSSA *S. aureus* ATCC 29213.

**Figure 6.**
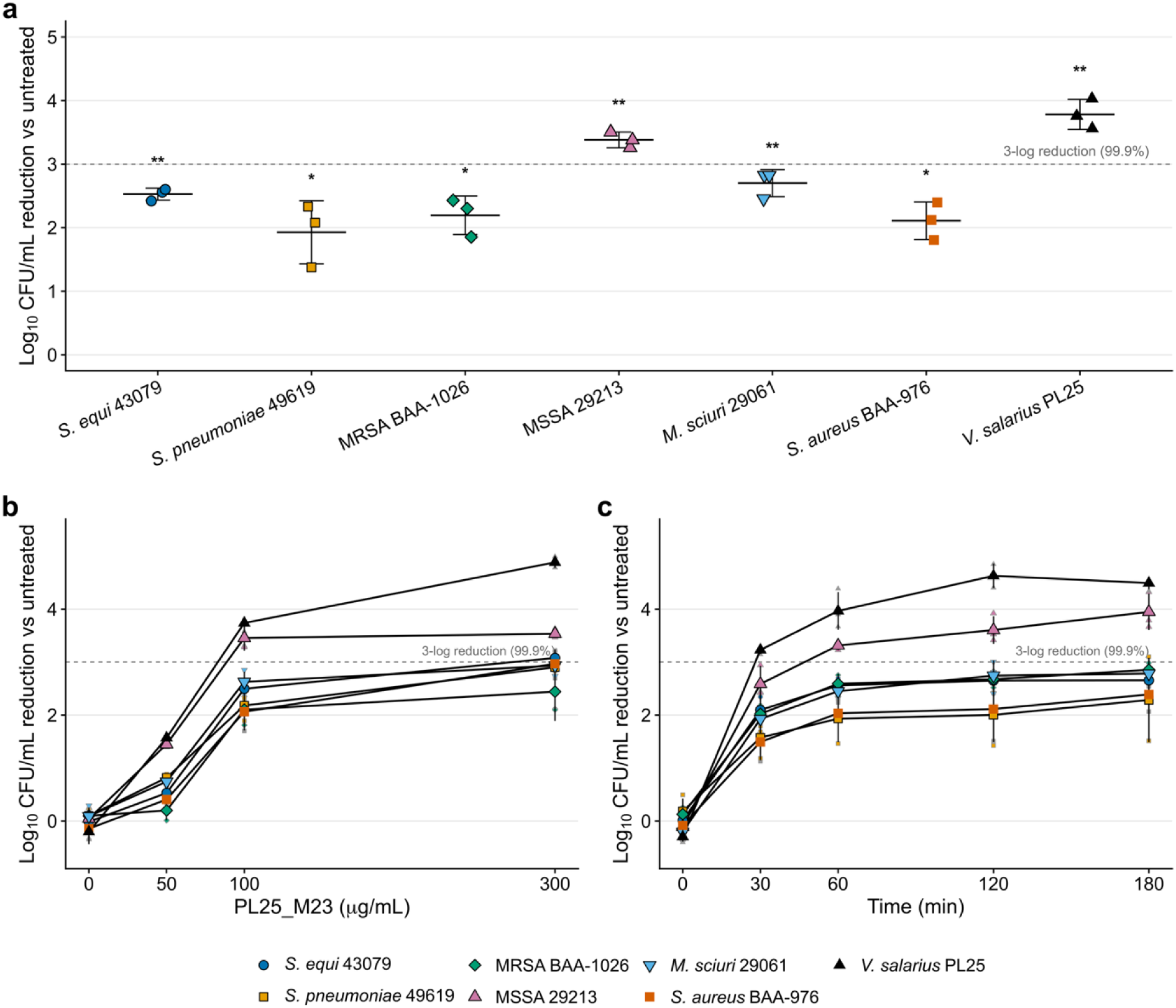
Concentration- and time-dependent antibacterial activity of PL25_M23 against Gram-positive bacteria. (a) Reduction in log10 CFU/mL relative to matched untreated controls following exposure to PL25_M23 (100 µg/mL, 60 min) across seven Gram-positive strains. Individual points represent independent biological replicates (*n* = 3), and horizontal lines indicate the mean ± SD. (b) Dose-response profiles following 60-min exposure to PL25_M23 at 0, 50, 100, or 300 µg/mL. (c) Time-kill kinetics following exposure to PL25_M23 at 100 µg/mL for 0–180 min. Values in panels (b) and (c) represent the mean ± SD of three independent biological replicates. Strain identity is indicated by marker shape and color, as shown in the legend. The dashed horizontal line indicates a 3-log10 reduction in viable counts (99.9%). Statistical comparisons were performed as described in Materials and Methods, and displayed P-values were adjusted using the Holm method. Storage-buffer controls, in which bacteria received the corresponding volume of protein storage buffer without enzyme, are shown in Supplementary Figure 2a.

Storage-buffer controls received the same volume of storage buffer as the corresponding PL25_M23 reactions. The resulting Δlog_10_ CFU/mL values remained close to zero at buffer carryover levels of 0%, 1.9%, and 3.8% (v/v), with mean values of +0.06, −0.01, and +0.04 log_10_ CFU/mL, respectively; none of these effects was statistically significant. At 11.5% (v/v), corresponding to the buffer carryover in the 300 µg/mL PL25_M23 treatment, the mean Δlog_10_ CFU/mL was +0.13 (adjusted P = 0.036). Thus, the largest buffer-associated effect was approximately 25-fold smaller than the 3.25-log_10_ reduction observed with 300 µg/mL PL25_M23. These results indicate that buffer carryover had a comparatively minor effect on viable counts and was unlikely to account for the antibacterial activity observed in the enzyme-treated samples (Supplementary Figure 2a).

### 3.7. PL25_M23 activity is attenuated by EDTA and increases under elevated salinity

The conserved HxxxD and HxH motifs in the predicted M23 domain prompted us to investigate whether PL25_M23 activity was sensitive to metal chelation. The addition of 1 mM EDTA attenuated PL25_M23-mediated killing across all seven strains, decreasing the mean reductions in viable counts from approximately 2.0–3.8 log_10_ CFU/mL with PL25_M23 alone to approximately 0.5–1.9 log_10_ CFU/mL in the presence of EDTA (Figure 7a). Supplementation with 1 mM ZnCl_2_ in the presence of EDTA partially restored antibacterial activity, yielding viable-count reductions of 1.4–3.6 log_10_ CFU/mL. The largest increases following ZnCl_2_ supplementation were observed for MRSA *S. aureus* ATCC BAA-1026 (from 1.1 to 2.4 log_10_ CFU/mL) and *V. salarius* PL25 (from 1.9 to 3.6 log_10_ CFU/mL). The addition of ZnCl_2_ to PL25_M23 in the absence of EDTA resulted in reductions of 2.3–3.8 log_10_ CFU/mL, comparable to those observed with PL25_M23 alone (1.9–3.8 log_10_ CFU/mL). Neither the EDTA-only nor the ZnCl_2_-only controls reproduced the multi-log reductions observed with the enzyme. Collectively, these findings support a contribution of metal availability to PL25_M23 activity but do not establish which catalytic domain is responsible for the observed antibacterial effect (Supplementary Figure 2b).

**Figure 7.**
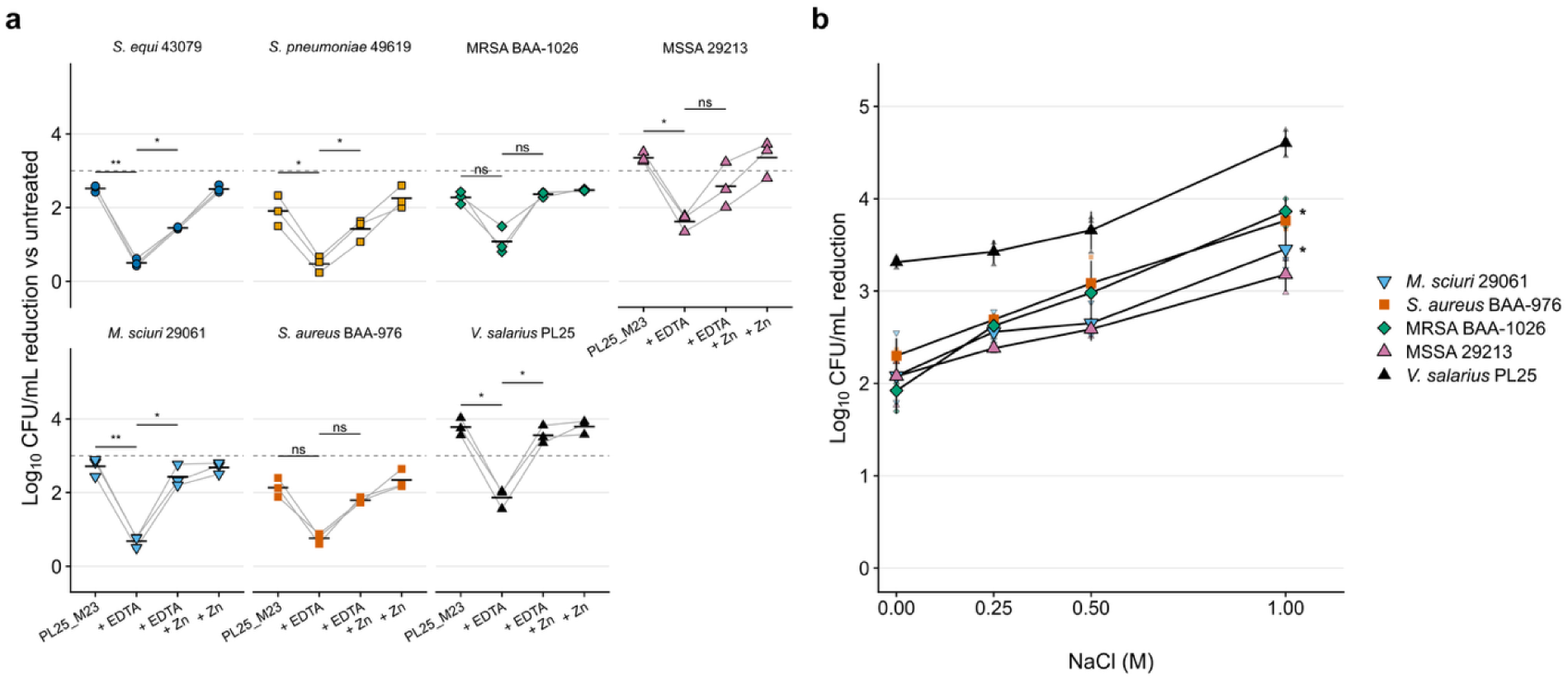
Metal sensitivity and PL25_M23 activity under elevated NaCl conditions. (a) Effects of EDTA and Zn^2+^ on the antibacterial activity of PL25_M23 across seven Gram-positive strains. Reductions in log10 CFU/mL relative to matched untreated controls are shown for PL25_M23 alone, PL25_M23 + 1 mM EDTA, PL25_M23 + 1 mM EDTA + 1 mM ZnCl2, and PL25_M23 + 1 mM ZnCl2. Values represent the mean ± SD of three independent biological replicates. Statistical brackets indicate the prespecified comparisons of PL25_M23 alone versus PL25_M23 + EDTA and PL25_M23 + EDTA versus PL25_M23 + EDTA + ZnCl2. P-values were adjusted using the Holm method, as described in Materials and Methods. EDTA-only and ZnCl2-only controls were included to assess the effects of these additives in the absence of PL25_M23. (b) PL25_M23 activity (100 µg/mL, 60 min) in standard PBS supplemented with 0, 0.25, 0.5, or 1.0 M additional NaCl, expressed as the reduction in log10 CFU/mL relative to the untreated control at 0 M added NaCl. Untreated controls were run at 0 and 1.0 M added NaCl only; values at 0.25 and 0.5 M are referenced to the 0-M untreated control. Five strains that retained viability at 1.0 M added NaCl were included. *S. equi* subsp. *zooepidemicus* ATCC 43079 and *S. pneumoniae* ATCC 49619 were excluded because 1.0 M additional NaCl alone reduced their viability by approximately 4.8 and 4.1 log10 CFU/mL, respectively. Values represent the mean ± SD of three independent biological replicates, with strain identity indicated by marker shape and color. Comparisons were performed within each strain against the 0-M additional NaCl condition using paired t-tests, with Holm correction across the 15 comparisons. Only statistically significant adjusted comparisons are annotated.

To evaluate PL25_M23 activity under elevated-salinity conditions, we tested the enzyme against the five strains that remained viable at 1.0 M added NaCl. *S. equi* subsp. zooepidemicus ATCC 43079 and *S. pneumoniae* ATCC 49619 were excluded because exposure to high salt alone reduced their viable counts by approximately 4.8 and 4.1 log_10_ CFU/mL, respectively (Supplementary Figure 2c). Among the five strains included in the analysis, the mean reductions in viable counts increased progressively with increasing NaCl concentration, from 2.34 log_10_ CFU/mL at 0 M added NaCl to 2.74, 2.99, and 3.77 log_10_ CFU/mL at 0.25, 0.5, and 1.0 M added NaCl, respectively (Figure 7b). Untreated controls were included at 0 and 1.0 M added NaCl; the 0.25 and 0.5 M values were therefore referenced to the 0-M untreated control rather than to salt-matched controls. Among the five retained strains, untreated viability did not differ significantly between 0 and 1.0 M added NaCl (mean difference, −0.05 log_10_ CFU/mL), supporting the use of the 0-M control as a common reference. The increase in viable-count reduction between 0 and 1.0 M NaCl ranged from 1.10 log_10_ CFU/mL for MSSA *S. aureus* ATCC 29213 to 1.94 log_10_ CFU/mL for MRSA *S. aureus* ATCC BAA-1026. Following Holm correction across the 15 comparisons, the 1.0 M condition differed significantly from the 0 M condition for MRSA *S. aureus* ATCC BAA-1026 and *M. sciuri* ATCC 29061. At 1.0 M added NaCl, for which salt-matched untreated controls were also available, comparisons of each enzyme-treated reaction with its corresponding salt-matched control showed increases in viable-count reduction of 1.20–1.71 log_10_ CFU/mL across all five strains; these differences were statistically significant in every strain. The greatest absolute reduction in viable counts was observed for the source strain, *V. salarius* PL25, reaching 4.60 log_10_ CFU/mL at 1.0 M added NaCl.

## 4. Discussion

### 4.1. PL25_M23 is associated with an under-characterized plasmid-candidate context

This study identified PL25_M23 as a divergent, predicted dual-domain SLT/M23 cell wall hydrolase encoded on pPL25-1 (contig_11), a 74-kb closed circular plasmid candidate identified in *V. salarius* PL25. Several independent observations support this genomic assignment, including circularity supported by junction-spanning Nanopore reads, higher sequence coverage than that of the main chromosomal contig, a high plasmid-classification score from geNomad, and a GC content distinct from that of the main chromosome (Camargo et al., 2024). Comparative analysis further showed that PL25_M23-related homologs occur in multiple halophilic or halotolerant Bacillaceae, including examples encoded on plasmids in closed genome assemblies. Collectively, these observations suggest that PL25_M23 belongs to an under-characterized group of Bacillaceae M23-family hydrolases associated with accessory genomic contexts (Vollmer, 2008; Vermassen et al., 2019). The mobility of pPL25-1, however, remains unresolved. Although the region contains genes annotated as plasmid maintenance- or transfer-associated markers, neither MOB-suite nor MacSyFinder/CONJScan identified a canonical MOB-family relaxase, a complete mating-pair formation system, or a complete conjugation module (Robertson and Nash, 2018; Cury et al., 2020). Accordingly, the available data do not establish that pPL25-1 is self-transmissible. Instead, the element may represent a divergent or incomplete plasmid, a non-mobilizable plasmid, or a plasmid that could be mobilized in trans. Experimental transfer assays and further characterization of its replication and transfer-associated regions will be needed to distinguish among these possibilities.

### 4.2. PL25_M23 has a CwlP-like predicted dual-domain SLT/M23 architecture

PL25_M23 combines two predicted catalytic modules within a compact 370-amino-acid protein: an N-terminal Slt35-/lysozyme-like glycan-hydrolase fold and a C-terminal Peptidase_M23 domain. This modular organization is consistent with the broader structural diversity of bacterial peptidoglycan hydrolases, in which distinct catalytic and substrate-targeting modules are frequently combined to support specialized cell wall remodeling functions (Vollmer, 2008; Vermassen et al., 2019). The closest architectural comparator identified in this study is CwlP, an SP-β prophage-associated hydrolase from *Bacillus subtilis* that contains an SLT-like muramidase domain and an M23 DD-endopeptidase domain (Sudiarta et al., 2010). Domain-restricted structural superposition showed a close correspondence between the predicted M23 domain of PL25_M23 and the M23 domain of CwlP, as well as broader structural similarity between their N-terminal glycan-hydrolase folds. The higher RMSD obtained in the unrestricted two-domain comparison suggests that, despite similarities in their individual catalytic-domain folds, PL25_M23 and CwlP may differ in their relative domain arrangements. Because AlphaFold2 models provide structural predictions rather than experimentally determined structures, these comparisons support a CwlP-like catalytic architecture but do not establish full-length protein homology or equivalent domain organization (Jumper et al., 2021). In contrast, CwlT combines a GH23-like bacterial lysozyme domain with an NlpC/P60 endopeptidase domain. Thus, PL25_M23 should not be considered CwlT-like at the domain-family level (DeWitt and Grossman, 2014; Xu et al., 2014). Moreover, domain annotation alone does not establish the precise reaction catalyzed by a protein. The N-terminal domain of CwlP was experimentally shown to possess muramidase activity despite its SLT-related annotation (Sudiarta et al., 2010). Accordingly, the predicted SLT-like fold of PL25_M23 should not be interpreted as evidence that it catalyzes a lytic transglycosylase reaction. The strain panel examined here spans two peptidoglycan chemotypes. Six of the seven strains possess A3α-type peptidoglycan, in which cross-linking involves an L-Lys-containing interpeptide bridge, whereas *V. salarius* PL25 possesses A1γ-type peptidoglycan, with direct cross-linking through *meso*-diaminopimelic acid (Schleifer and Kandler, 1972; Kloos et al., 1976; Hua et al., 2008). PL25_M23 reduced viable counts in strains representing both chemotypes. This activity is noteworthy because the substrate specificities of characterized endopeptidases such as lysostaphin and the CwlP M23 domain are associated with particular peptidoglycan cross-link structures. Lysostaphin cleaves the glycyl-glycine bonds in the interpeptide bridges of susceptible staphylococci, whereas the CwlP M23 domain has been reported to cleave DAP-containing peptidoglycan but not the peptidoglycan of Staphylococcus aureus or Streptococcus thermophilus (Sudiarta et al., 2010). The activity of PL25_M23 against both chemotypes is therefore compatible with either a peptidase with broader substrate specificity or a contribution from its predicted N-terminal glycan-hydrolase domain, which may act on a peptidoglycan component shared by both chemotypes. The present experiments do not distinguish between these possibilities. Although residual killing in the presence of EDTA ranged from 20% to 49% of the uninhibited reduction, this residual activity did not show an apparent separation according to peptidoglycan chemotype. Neither catalytic activity has yet been demonstrated directly for PL25_M23. Because viable-count reductions do not identify the peptidoglycan bonds cleaved, establishing whether PL25_M23 has dual catalytic activity will require assays of separately expressed domains and direct analysis of peptidoglycan cleavage products.

### 4.3. PL25_M23 combines an acidic surface with salt-enhanced and metal-sensitive activity

PL25_M23 exhibited a more acidic predicted surface profile than the reference hydrolases examined in this study. Its modeled catalytic region had a low predicted pI, a negative net charge at pH 7.0, and acidic side chains distributed across the predicted protein surface. These characteristics are consistent with physicochemical features commonly reported for halophilic and halotolerant proteins. In such proteins, enrichment in acidic residues and negatively charged surface patches may contribute to hydration, solubility, and structural stability under conditions of high ionic strength (Madern et al., 2000; Oren, 2008; Paul et al., 2008; Graziano and Merlino, 2014). Nevertheless, an acidic surface profile is not, by itself, evidence of adaptation to high salinity. PL25_M23 activity was attenuated by EDTA across all seven strains, while the addition of ZnCl_2_ partially restored antibacterial activity. These findings support a contribution of metal availability to PL25_M23 activity and are compatible with the conserved HxxxD and HxH motifs in its predicted M23 domain, which are characteristic of the metal-binding architecture of many M23-family peptidases. The contrast with the Zn^2+^-independent M23 domain of CwlP also illustrates that proteins sharing an M23 domain architecture do not necessarily have identical metal requirements. However, EDTA can chelate multiple divalent cations, and Zn^2+^ was the only cation tested for restoration of activity. The experiments therefore support metal sensitivity and show that Zn^2+^ supplementation can restore activity under the conditions tested, but they do not establish Zn^2+^ as the physiological cofactor or exclude the possibility that other cations, such as Ca^2+^, Mg^2+^, or Mn^2+^, could also support activity. Furthermore, the whole-protein viable-count assay does not demonstrate that EDTA acts exclusively through the M23 domain. The second predicted catalytic domain, which has a glycan-hydrolase-like fold, could contribute to the residual antibacterial activity observed after EDTA treatment. Resolving the contribution of each domain will require catalytic-site mutagenesis, assays of isolated domains, metal-content measurements, and direct analysis of peptidoglycan cleavage products. Elevated NaCl concentrations increased rather than decreased the antibacterial activity of PL25_M23 in every strain included in the salinity experiment. Across the five strains tested, the mean reduction in viable counts increased by approximately 1.4 log_10_ CFU/mL between 0 and 1.0 M added NaCl. This enhancement is consistent with adaptation to a saline environment, in which elevated salt may support protein activity rather than merely being tolerated (Madern et al., 2000), and with the unusually acidic predicted surface profile of PL25_M23. A comparable response has been reported for lysostaphin: although NaCl inhibits its catalytic activity and cell-wall binding under some assay conditions, it can increase the bacteriolytic activity of the full-length enzyme, an effect attributed to changes in peptidoglycan conformation rather than to a direct stimulatory effect on the enzyme (Konstantinova et al., 2022). Two non-exclusive mechanisms could account for the salt-associated increase in PL25_M23 activity. First, the predicted net charge of −22.4 at pH 7.0 may impede the approach of PL25_M23 to the negatively charged, teichoic-acid-rich Gram-positive cell envelope through electrostatic repulsion (Neuhaus and Baddiley, 2003; Brown et al., 2013). Increasing the ionic strength of the medium could screen these electrostatic interactions and facilitate enzyme– cell-wall association (Schreiber et al., 2009). Second, added NaCl may alter the conformation or accessibility of the peptidoglycan substrate, as proposed for the salt-associated increase in lysostaphin activity. Notably, the salt tolerance of lysostaphin has been linked to its cell-wall-binding domain (Jagielska et al., 2016), whereas PL25_M23 has no recognizable cell-wall-binding module yet exhibited increased activity at elevated NaCl concentrations. Distinguishing between effects on enzyme–cell-wall association and effects on substrate accessibility will require direct binding and activity measurements across a range of ionic strengths, together with analyses of charge-reversal variants.

### 4.4. Reproducible antibacterial activity supports further enzybiotic evaluation

Recombinant PL25_M23 reduced viable counts across seven Gram-positive strains representing four genera, with activity observed in three independent biological experiments. The pronounced increase in activity between 50 and 100 µg/mL, together with the rapid onset of killing, suggests a concentration-dependent response with substantial activity developing over a relatively narrow concentration range. Although most of the reduction occurred within the first 30–60 min, additional decreases in viable counts were observed at later time points in several strains. Activity against both MRSA and MSSA also extends the potential relevance of PL25_M23 beyond its source strain, *V. salarius* PL25. The observation that PL25_M23 produced its largest reduction in viable counts against *V. salarius* PL25 itself is noteworthy because the gene is encoded on a replicon identified in this strain. However, the assay involved applying purified enzyme externally to washed bacterial cells, and this experimental configuration does not necessarily reflect the accessibility or physiological role of PL25_M23 at the cell surface in its native host. More broadly, the observed activity is consistent with previous studies demonstrating that peptidoglycan hydrolases, including phage endolysins and lysostaphin-family enzymes, can act rapidly against Gram-positive bacteria (Nelson et al., 2001; Fischetti, 2010; Schmelcher et al., 2012; Sabala et al., 2014). These findings establish reproducible in vitro antibacterial activity but do not yet constitute therapeutic validation. Further work should include independent protein-production batches, testing against a broader range of bacterial strains, direct characterization of peptidoglycan cleavage, cytotoxicity assessment, and evaluation of resistance development before PL25_M23 can be considered a validated enzybiotic candidate. An additional limitation concerns the preparation used in this study: the recombinant protein was recovered from inclusion bodies, and the proportion of correctly folded, enzymatically active protein in the final preparation is unknown. Consequently, activity normalized to total protein concentration may underestimate the specific activity of the correctly folded fraction. Although the attenuation by EDTA and partial restoration by Zn^2+^ are consistent with the presence of a metal-responsive active fraction, these results do not establish the proportion of active protein in the preparation. Quantifying this fraction will require a soluble, appropriately folded preparation and a direct assay of peptidoglycan-cleavage activity.

## 5. Conclusion

This study identified PL25_M23 as a compact, plasmid-candidate-associated, predicted dual-domain SLT/M23 cell wall hydrolase from the halotolerant bacterium *V. salarius* PL25. Genomic, phylogenetic, and structural analyses support its placement among under-characterized Bacillaceae M23-family hydrolases and suggest a CwlP-like catalytic architecture. Recombinant PL25_M23 exhibited reproducible, concentration- and time-dependent antibacterial activity against seven Gram-positive strains. Its activity was sensitive to metal chelation and was partially restored by Zn^2+^ supplementation, while elevated salinity increased its antibacterial activity. These findings establish PL25_M23 as a candidate for further mechanistic investigation and enzybiotic development. However, plasmid mobility, the catalytic activities of its individual domains, and its therapeutic potential remain to be established experimentally.

## Data availability

The genome assembly and annotation of *V. salarius* PL25 have been deposited in DDBJ/ENA/GenBank under BioProject PRJNA1473924 and BioSample SAMN60541943, with the locus-tag prefix AC3VG8. The Whole Genome Shotgun project has been deposited under accession JBYTNQ000000000; version JBYTNQ010000000 will be released upon acceptance of this manuscript. Raw Oxford Nanopore reads are available from the NCBI Sequence Read Archive under accession SRR35929825. The PL25 isolate is also associated with the umbrella BioProject PRJNA1356516, “Pink Lake Isolate from Allith, Saudi Arabia.” Source data for Figures 1–7 are provided in Supplementary Data S1. Analysis scripts and intermediate files are available from the corresponding author upon reasonable request.

## Author contribution

**A.H. Farran** contributed to conceptualization, methodology, investigation, visualization, and writing of the original draft. **N. Kontis** contributed to data curation, resources, and writing (review and editing). **A. Rosado** contributed to supervision, project administration, funding acquisition, and writing (review and editing). All authors contributed to the article and approved the submitted version.

## Acknowledgments

We thank Prof. Danesh Moradigaravand and members of the Moradigaravand laboratory for generously providing the ATCC reference strains used in this study.

## Supplementary material

The Supplementary Material for this article will be provided as part of the manuscript submission.

