## Supplementary Methods for "PL25_M23: Salt-enhanced antibacterial activity of a plasmid-encoded SLT–M23 hydrolase from *Virgibacillus salarius* against Gram-positive bacteria"

**Methods S1: Confirmation of contig_11 circularity from the Nanopore reads**

Flye reports contig_11 as circular but trims the closure overlap before writing the final contig, so a circular and a linear contig cannot be distinguished at the FASTA level. Circularity was therefore tested directly from the reads. Two 40-kb references were built from contig_11 (74,025 bp): a junction reference consisting of the final 20 kb joined to the first 20 kb, placing the assembler's end-to-start join at position 20,000; and an internal control consisting of an interior 40-kb window with its midpoint at the same position. The Oxford Nanopore reads were mapped to both references in a single pass with minimap2 (-ax map-ont), unmapped reads were discarded, and alignments were sorted and indexed with SAMtools. A read was counted as spanning only if at least 1 kb aligned on both sides of position 20,000; secondary and supplementary alignments were excluded so that a read split across the join could not be counted as spanning it. The internal control provides the spanning-read rate expected for genuinely contiguous sequence at the same coverage and read-length distribution.

**Methods S2: CONJScan/MacSyFinder Command**

Conjugation-system completeness was assessed using MacSyFinder v2.1.6 and the CONJScan v2.1.0 plasmid model set:

macsyfinder \

--models CONJScan/Plasmids all \

--db-type ordered_replicon \

--replicon-topology circular \

--hmmer hmmsearch \

-w 4 \

--sequence-db results/03_M23_ICE_locus/contig_11_proteome.faa \

-o results/17_conjscan/contig11_run

### The CONJScan/Plasmids all option screened the 99 predicted proteins of contig_11 against all plasmid conjugation models in CONJScan v2.1.0. No complete conjugation system was detected; partial hits are summarized in **Supplementary Table S1**

### **Methods S3: geNomad Command Line and oriT motif sequences, primary references, and scan parameters**

### geNomad v1.7.0 was run in end-to-end mode against the geNomad v1.7 database:

### genomad end-to-end \

### --threads 8 --cleanup --splits 4 \

### --min-score 0.7 \

### --min-plasmid-marker-enrichment 0 \

### --min-virus-marker-enrichment 0 \

### --min-plasmid-hallmarks 0 \

### --min-virus-hallmarks 0 \

### PL25_working.fna results/06_genomad/ /path/to/genomad_db

### Marker-enrichment thresholds were relaxed to improve detection of divergent plasmid-like elements. Per-contig scores are provided in **Supplementary Table S3.**

Five canonical *nic*-site core motifs were tested against both strands of PL25 contig_11 (74,025 bp). Motif sequences below are the consensus published in each primary reference; ambiguity codes follow IUPAC.

| Motif | Consensus sequence (5′→3′) | Primary reference |
| --- | --- | --- |
| IncP RP4 | CTGCCTGGTGCT | Pansegrau & Lanka, 1991 |
| IncQ RSF1010 | GGGTGYGNGCGRCY (MobA recognition) | Becker & Meyer, 2000 |
| IncW R388 | TTGTTGTRYRGTRCRYR (TrwC inverted repeat + loop) | Llosa et al., 1994 |
| pIP501 (Gram+) | TAGTGCGCC (orfH/orfJ region) | Wang & Macrina, 1995 |
| ICEBs1 nic-site core | CTTGTCTAGTTATAA (NicK recognition) | Lee et al., 2010 |

A hit was scored as positive only if it fell within **± 3 kb of the predicted relaxase candidate PL25_00076** and was flanked by an **inverted repeat of ≥ 18 bp** (stride 1 bp, up to two mismatches in the stem). sequence-level matches. Results are summarized in **Supplementary Table S1.**

### **Methods S4: Pfam HMM panel (12 profiles)**

The HMM panel of **12 Pfam profiles** scanned against the Prokka proteome of PL25 contig_11 with HMMER v3.4 hmmscan at E ≤ 10^-5^, using Pfam release 37.0

| Category | Pfam ID | Profile name | Function |
| --- | --- | --- | --- |
| **Cell-wall hydrolase (M23)** | PF01551 | Peptidase_M23 | Zn^2+^-dependent endopeptidase |
| **Cell-wall hydrolase (SLT)** | PF01464 | SLT | Soluble lytic transglycosylase |
| **Cell-wall hydrolase (GH25)** | PF01183 | Glyco_hydro_25 | GH25 muramidase fold |
| **Cell-wall hydrolase (NlpC/P60)** | PF00877 | NLPC_P60 | NlpC/P60 cysteine peptidase |
| **Relaxase (replication initiator)** | PF03389 | MobA_MobL | Plasmid replication-relaxation |
| **T4SS** | PF02534 | T4SS_TraD-like | DNA-transfer coupling protein |
| **VirB4-like ATPase** | PF03135 | TrwB_AAD_bind | VirB4-family AAA-ATPase |
| **AAA-domain ATPases** | PF13245 | AAA_19 | RecD2-like helicase (matches PL25_00094) |
|  | PF13191 | AAA_16 | Generic AAA ATPase |
|  | PF12846 | AAA_10 | AAA-domain ATPase, VirB4-associated |
| **DNA-repair** | PF05866 | RusA | Holliday-junction resolvase |
| **Cell-wall-binding** | PF08239 | SH3_3 | SH3b cell-wall-targeting domain |

### **Methods S5: PyMOL**

Surface representations of PL25_M23 and the four reference proteins (CwlP, CwlT, lysostaphin, LytM) used in Figure 5a were generated in PyMOL v3.1.6.1 (Open-Source) using /Applications/PyMOL.app/Contents/bin/pymol -cq scripts/08_render_surfaces.py. A **single per-atom side-chain coloring rule** was applied identically to every protein so that the resulting surfaces are comparable:

| Side chain | Color | Rationale (at pH 7) |
| --- | --- | --- |
| Asp (D), Glu (E) | red | acidic (~ –1) |
| Lys (K), Arg (R) | blue | basic (~ +1) |
| His (H) | pink | partial protonation |
| All other residues | white / neutral | uncharged |

The PyMOL commands used were:

cmd.load(pdb_path, name)
cmd.show_as("surface", name)
cmd.color("white", name)
cmd.color("red", f"({name}) and resn ASP+GLU and (sidechain or name CA)")
cmd.color("blue", f"({name}) and resn LYS+ARG and (sidechain or name CA)")
cmd.color("pink", f"({name}) and resn HIS and (sidechain or name CA)")

Front and 180° rotated views were captured with cmd.rotate("y", 180) between renders. Images were ray-traced at width 1200 × height 1200 (cmd.ray(1200, 1200)) before saving as PNG. PL25_M23 (residues 38–370) and the CwlP / CwlT comparators were rendered from the AlphaFold2 models; lysostaphin (PDB 4QPB chain A, mature) and LytM (PDB 4ZYB chain A, mature) were rendered from the experimental crystal structures.
