## Supplementary Tables and Figs for "PL25_M23: Salt-enhanced antibacterial activity of a plasmid-encoded SLT–M23 hydrolase from *Virgibacillus salarius* against Gram-positive bacteria"

**Table S1:**

| Test | Tool | Reference profiles | Threshold | Result on contig_11 |
| --- | --- | --- | --- | --- |
| **Plasmid replicon typing** | MOB-suite v3.1.9 | rep_clusters database (Robertson & Nash 2018) | Closest cluster within Mash distance ≤ 0.05 | **No replicon detected** (closest hit: *Spiroplasma citri* CP042474 at Mash distance 0.296, beyond the typing threshold) |
| **Relaxase typing** | MOB-suite v3.1.9 | MOB family HMMs (MOBP / F / V / Q / C / H) | E ≤ 10^-30^ to any canonical MOB profile | **None of the six MOB families detected** |
| **Mating pair formation (MPF) typing** | MOB-suite v3.1.9 | MPF_T / F / G / I / B / C / FATA profiles | Standard MPF cluster assignment | **No MPF type** |
| **oriT typing** | MOB-suite v3.1.9 | oriT consensus library | Match to known oriT sequence | **No oriT detected** |
| **Conjugation system completeness** | MacSyFinder v2.1.6 + CONJScan v2.1.0 (Cury et al. 2020) | 17 plasmid models: MOB, T4SS_typeB/C/F/FA/FATA/G/I/T, and decay (dCONJ_*) variants | Mandatory-gene quorum reached per model | **0 complete systems detected** across all 17 models. Partial / subquorum hits only: PL25_00148 → T4SS coupling protein (T4SS_t4cp2, HMM score 77); PL25_00150 → VirB11-like ATPase (T4SS_T_virB11, HMM score 79); PL25_00085 → FA_orf13 (weak, score 15). **No MOB-relaxase hit** at any threshold |
| **Canonical oriT motif scan** | Local Python scan | IncP RP4 (CTGCCTGGTGCT), IncQ RSF1010, IncW R388, pIP501 Gram+ (TAGTGCGCC), ICEBs1 nic-site core | motif match within ± 3 kb of relaxase candidate + flanking long inverted repeat | **Negative** for all motifs adjacent to the relaxase candidate (PL25_00076). Two level ICEBs1 13-mer matches detected ≥ 24 kb away (positions 1,171 and 30,570 of 74,025 bp) with no inverted-repeat. No long (≥ 18 bp) inverted repeat near the relaxase |

### **Table S2:**

**Structural superposition of PL25_M23 against ref M23 and cell-wall hydrolases.** All superpositions were performed in PyMOL v3.1.6.1 using cmd.super() with five cycles of outlier rejection at 2.0 Å. The query structure was the AlphaFold2 *ptm* model of PL25_M23 (residues 38–370 of the full-length 370-aa protein; mean pLDDT = 88.7; ColabFold v1.5.5). Reference structures were obtained from the RCSB PDB or from the AlphaFold Protein Structure Database (accessed 2026-05-22). Atom counts are Cα pairs retained after PyMOL's outlier rejection. Residue numbering throughout this table refers to **full-length PL25_M23** (consistent Figures 3, 4 and 5). Values are in Figure 4 (PL25 alone) and 5 (PL25 vs CwlP vs CwlT).

#### **A. C-terminal peptidase domain**

| Reference | Source organism / location | Structure used | Peptidase family | RMSD (Å) | Aligned residues |
| --- | --- | --- | --- | --- | --- |
| **CwlP (M23)** | *B. subtilis* SP-β prophage, catalytic core res 1373-1686 of UniProt O31976 | AlphaFold2 model (ColabFold v1.5.5) | M23 (PF01551) | **0.69** | **117** |
| Lysostaphin | *Staphylococcus simulans*, secreted enzybiotic | PDB 4QPB chain A | M23 (PF01551) | 0.49 | 102 |
| LytM | *Staphylococcus aureus*, autolysin | PDB 4ZYB chain A | M23 (PF01551) | 0.60 | 115 |
| ALE-1 | *Staphylococcus capitis*, secreted lysin | AlphaFold2 model AF-O05156-F1 v6^1^ | M23 (PF01551) | 0.42 | 80 |
| Zoocin A | *Streptococcus equi* subsp. *zooepidemicus* | PDB 5KVP | M23 (PF01551) | 1.03 | 90 |
| CwlT (C-terminal peptidase) | *B. subtilis* ICE *ICEBs1* | AlphaFold2 model AF-P96645-F1 v4 | **NlpC/P60 (PF00877)** different family | **3.23** | **16** |

#### **B. N-terminal glycan-hydrolase domain**

| Reference | Source organism / location | Structure used | N-terminal-domain family / fold | RMSD (Å) | Aligned residues |
| --- | --- | --- | --- | --- | --- |
| **CwlP-clipped (SLT)** | *B. subtilis* SP-β prophage, SLT domain | AlphaFold2 model (ColabFold v1.5.5) | SLT / Lysozyme-like (PF01464) | **3.31** | **108** |
| CwlT (N-terminal glycan-hydrolase) | *B. subtilis* ICE *ICEBs1* | AlphaFold2 model AF-P96645-F1 v4 | **GH25 muramidase (PF01183)** distinct fold | 10.43 | 109 |

#### **C. Predicted two domain regions**

PL25_M23 residues 38-370 (the modelled region) were superposed onto each reference enzyme without domain restriction.

| Comparison | Architecture | RMSD (Å) | Aligned residues |
| --- | --- | --- | --- |
| **PL25_M23 vs CwlP-clipped** | SLT + M23 (same architecture) | **16.75** | **234** |
| PL25_M23 vs CwlT | GH25 + NlpC/P60 (different family) | 10.43 | 109 |

The higher RMSD obtained for the CwlP comparison under C table, relative to table A and table B, indicates that the catalytic domain folds of PL25_M23 and CwlP are conserved, but their inter domain hinge orientation has diverged. The CwlT comparison is included in panel C only to illustrate the architectural mismatch, different catalytic families

Notes

1. ALE-1: PDB 1R77 covers only the C-terminal SH3b domain (residues 271-362). The AlphaFold2 monomer AF-O05156-F1 v6 was used here because it includes the M23 catalytic core.
2. CwlT identity: *B. subtilis* CwlT (UniProt P96645) is a GH25 + NlpC/P60 hydrolase, not M23. PDB 4FDY in fact maps to UniProt Q932I6 (a *S. aureus* NlpC/P60 lipoprotein). The AlphaFold2 model of the correct P96645 sequence is used here, and CwlT is included only as an architectural negative control.
3. AF2-vs-AF2 caveat: no experimental structures exist for CwlP (O31976) or *B. subtilis* CwlT (P96645), the PL25 vs CwlP and PL25 vs CwlT comparisons are therefore AlphaFold2 vs AlphaFold2, whereas PL25 vs lysostaphin/LytM/EnpA/zoocin A are AlphaFold2 vs crystal.

### **Table S3:**

**Flye reported circularity, and geNomad classification of PL25 contigs.** Throughout the manuscript and this table, **contig_11** refers to the 74,025-bp plasmid candidate that carries PL25_M23 (locus tag PL25_00078).

| Manuscript contig (FASTA) | Flye contig | Length (bp) | n CDS | Flye circ | Flye cov. (×) | Flye mult. | GC % | GC3 % | ΔGC vs chr. | ΔGC3 vs chr. | geNomad plasmid score | geNomad virus score | Final assignment |
| --- | --- | --- | --- | --- | --- | --- | --- | --- | --- | --- | --- | --- | --- |
| **contig_4** | contig_4 | 4,349,560 | 4,098 | N | 476 | 1 | 36.81 | 28.71 | - | - | 0.097 | - | chromosomal contig |
| contig_2 | contig_2 | 249,289 | - | N | 849 | 2 | 36.29 | - | -0.52 | - | - | - | Unannotated chromosomal fragment (no Prokka CDS, high coverage and elevated multiplicity consistent with repeat) |
| **contig_11** | **contig_12** | **74,025** | **99** | **Y** | **889** | **2** | **31.22** | **23.19** | **-5.59** | **-5.52** | **0.989** | - | **Closed circular plasmid candidate (contains PL25_M23)** |
| contig_12 | contig_11 | 44,498 | 86 | Y | 22 | 1 (alt 2) | 36.31 | 30.85 | -0.50 | +2.14 | - | 0.979 | Closed circular, repeat-flagged; geNomad virus-classified (likely a free prophage) |
| contig_10 | contig_10 | 32,390 | 70 | Y | 86 | 1 | 29.70 | 20.59 | -7.11 | -8.12 | 0.967 | - | Closed circular replicon |
| contig_9 | contig_9 | 3,944 | 5 | N | 4 | 1 | 35.45 | 26.99 | -1.36 | -1.72 | 0.818 | - | Short linear fragment |
| contig_6 | contig_6 | 514 | 0 | N | 3 | 1 | 40.86 | - | +4.05 | - | - | - | Short linear fragment (< 600 bp, no Prokka CDS) |
| contig_5 | contig_5 | 462 | 0 | N | 4 | 4 | 41.34 | - | +4.53 | - | - | - | Short linear fragment (< 600 bp, no Prokka CDS) |

| Flye contig | Length (bp) | Flye circ | Flye cov. (×) | Reason for exclusion |
| --- | --- | --- | --- | --- |
| contig_1 | 3,815 | N | 4 | Failed length (< 5 kb) and coverage (< 10×) thresholds. Excluded |

### Notes

### Flye circ = Y is taken from assembly_info.txt. For contig_11 (pPL25-1), circularity was additionally verified directly from the Nanopore reads by a junction-spanning read test (Supplementary Methods S1; Results). For contig_12 and contig_10, Flye circularity was not independently verified.

### Δ-values are differences vs the chromosomal baseline (contig_4: GC 36.81 %, GC3 28.71 %).

### ΔGC3 may indicate horizontal acquisition (Lawrence & Ochman, 1998), but must be read together with the conjugation/synteny analyses (supplementary Table S1, Figure 1b)

Contig identifiers were reassigned by length in the final assembly, so the two largest extrachromosomal contigs are interchanged relative to the original Flye output: manuscript contig_11 (pPL25-1, 74,025 bp) is Flye contig_12, and manuscript contig_12 (44,498 bp) is Flye contig_11. All other contigs retain their Flye identifiers. The “Flye contig” column gives the original identifier for each row

### **Table S4:**

**PL25_M23 homologues in closed NCBI Bacillaceae assemblies: replicon context, protein hit, and BLASTp statistics.**

| **rank** | **organism** | **assembly** | **replicon** | **plasmid_name** | **nucleotide_acc** | **protein_acc** | **length_aa** | **pct_identity** | **evalue** |
| --- | --- | --- | --- | --- | --- | --- | --- | --- | --- |
| **1** | Cytobacillus oceanisediminis 2691 | GCF_000294775.2 | ★ plasmid | pBO1 | NZ_CP015507.1 | WP_019380455.1 | 703 | 44.8% | 4.7 × 10^-65^ |
| **2** | Halobacillus litoralis ERB 031 | GCF_004101865.1 | ★ plasmid | pLDW-31 | NZ_CP026119.1 | WP_241656025.1 | 620 | 43.8% | 1.8 × 10^-61^ |
| **3** | Virgibacillus halodenitrificans Bac324 | GCF_003667805.1 | ★ plasmid | unnamed | NZ_CP033050.1 | WP_121616750.1 | 758 | 48.4% | 3.8 × 10^-65^ |
| **4** | Gracilibacillus boraciitolerans JCM 21714 | GCF_055385745.1 | ★ plasmid | pJCM21714 | NZ_AP043914.1 | WP_464380148.1 | 363 | 50.0% | 1.1 × 10^-106^ |
| **5** | Salinicoccus jeotgali JCM 16981 | GCF_055379195.1 | ★ plasmid | pJCM16981-2 | NZ_AP043832.1 | WP_344700570.1 | 426 | 35.0% | 6.7 × 10^-61^ |
| **6** | Cytobacillus firmus M7 | GCF_025732135.1 | ★ plasmid | p1 | NZ_CP107028.1 | WP_263600221.1 | 707 | 44.5% | 2.7 × 10^-64^ |
| **7** | Cytobacillus firmus CK19 | GCF_023823815.1 | ★ plasmid | unnamed1 | NZ_CP085391.1 | WP_252222728.1 | 707 | 44.5% | 2.9 × 10^-64^ |
| **8** | Cytobacillus firmus SGAir0285 | GCF_043540845.1 | ★ plasmid | pSGAir0285_3 | NZ_CP027985.1 | WP_009335593.1 | 627 | 43.4% | 1.7 × 10^-61^ |
| **9** | Cytobacillus oceanisediminis CK22 | GCF_023823835.1 | ★ plasmid | unnamed1 | NZ_CP085393.1 | WP_252246573.1 | 707 | 44.5% | 3.6 × 10^-64^ |
| **10** | Cytobacillus oceanisediminis YPW-V2 | GCF_014883935.1 | ★ plasmid | unnamed1 | NZ_CP062791.1 | WP_192909257.1 | 627 | 43.8% | 2.4 × 10^-61^ |
| **11** | Cytobacillus pseudoceanisediminis | GCF_023516215.1 | ★ plasmid | unnamed | NZ_CP097350.1 | WP_251264864.1 | 707 | 44.5% | 2.3 × 10^-63^ |
| **12** | Lentibacillus sp. CBA3610 | GCF_013373365.1 | chromosome | — | NZ_CP035925.1 | WP_176447356.1 | 361 | 52.9% | 8.2 × 10^-128^ |
| **13** | Lentibacillus amyloliquefaciens LAM0015 | GCF_001307805.1 | chromosome | — | NZ_CP013862.1 | WP_068441797.1 | 358 | 51.1% | 6.7 × 10^-123^ |
| **14** | Virgibacillus ainsalahensis JCM 30907 | GCF_055380755.1 | chromosome | — | NZ_AP043943.1 | WP_461177648.1 | 360 | 53.0% | 6.9 × 10^-126^ |

#### Notes

1. **Multi-domain plasmid-encoded homologues**: plasmid-encoded homologues range from 363 aa to 758 aa, with most (n = 9) in the 620–758 aa range, compared with chromosomal homologues (358–361 aa) and PL25_M23 itself (370 aa). The larger plasmid-encoded variants are likely multi-domain fusion proteins; the identity of their additional N-/C-terminal domains was not investigated in this study.
2. **% identity and E-value** are taken from the original BLASTp search of PL25_M23 (38–370 aa) against NCBI nr (-evalue 1e-5 -max_target_seqs 250); see main Methods. Identity is to the full-length BLAST alignment region, not to the PL25_M23 M23 catalytic domain alone.

### **Table S5:**

**M23 Zn^2+^-binding HxxxD + HxH motifs on PL25_M23**

Residues were identified by sequence motif scan of each parent structure / model and verified by visual inspection of the M23 catalytic site after pairwise superposition onto PL25_M23 (PyMOL v3.1.6.1, cmd.super, five outlier-rejection cycles at 2.0 Å). M23-domain RMSDs (Cα) versus PL25_M23 residues 242-367 are listed for context; all four reference enzymes superpose into a narrow 0.42-0.69 Å band, confirming a structurally conserved catalytic core. Residue positions for PL25_M23 refer to the **full-length 370-aa sequence**, Figure 3.

| Protein (parent structure) | Source | HxxxD: H1 | HxxxD: D | HxH: H2 | HxH: H3 |
| --- | --- | --- | --- | --- | --- |
| **PL25_M23** | *V. salarius* PL25 plasmid contig_11 | **H264** | **D268** | **H346** | **H348** |
| Lysostaphin (PDB 4QPB) | *Staphylococcus simulans*, secreted enzybiotic | H279 | D283 | H360 | H362 |
| LytM (PDB 4ZYB) | *Staphylococcus aureus*, autolysin | H210 | D214 | H291 | H293 |
| ALE-1 (AF2 of UniProt O05156) | *Staphylococcus capitis*, secreted lysin | H150 | D154 | H231 | H233 |
| CwlP-clipped (AF2 of UniProt O31976, residues 1373-1686) | *Bacillus subtilis*, SP-β prophage; reported Zn^2+^-independent | H1580 (H208 of clip) | D1584 (D212) | H1660 (H288) | H1662 (H290) |

Notes

1. ALE-1: PDB 1R77 covers only the C-terminal SH3b domain (residues 271-362); AlphaFold2 monomer AF-O05156-F1 v6 was used here because it includes the M23 catalytic core.
2. CwlP-clipped numbering: full-length CwlP is 2,285 aa (SP-β prophage tail-associated protein); the SLT + M23 catalytic core (residues 1373-1686 of UniProt O31976) was modelled with ColabFold v1.5.5 (AlphaFold2 ptm). Residue numbers in the table give both the full-length position and the equivalent in the 314-aa clipped construct.
3. CwlP Zn^2+^-independence: Sudiarta et al. (2010, *J. Biol. Chem.* 285:41232- 41243) reported that CwlP's M23 domain does not require Zn^2+^ for activity, yet our motif scan shows CwlP retains both canonical motifs (H1580–D1584 and H1660/H1662). Motif conservation therefore does not establish Zn^2+^ dependence; confirmed (EDTA inhibition, Zn^2+^ titration).
4. CwlT omitted: *B. subtilis* CwlT (UniProt P96645) is a GH25 + NlpC/P60 hydrolase and carries no M23 catalytic domain, so it is excluded from this motif-conservation table. See Supplementary Table S2 note 3 for the PDB 4FDY.
5. Motif scan: in-house Python regex H[A-Z]{3}D (HxxxD) and H[A-Z]H (HxH), applied to the primary sequence of each parent structure within the predicted M23 catalytic domain.

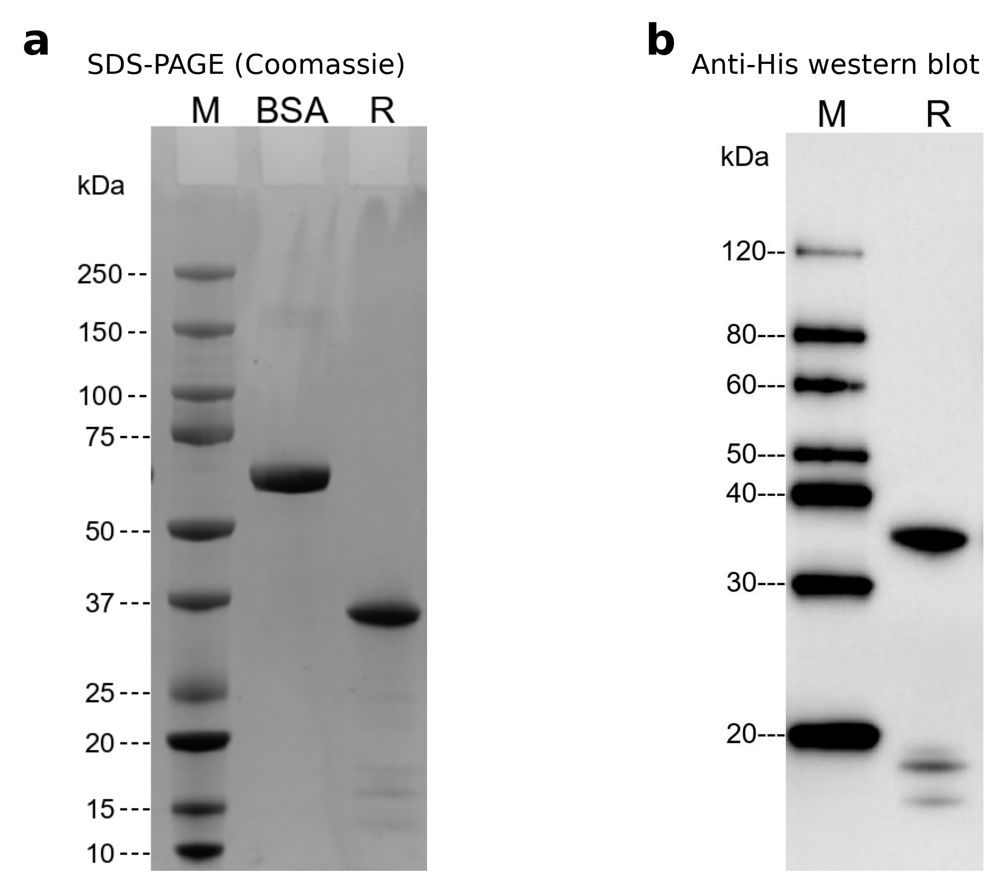

**Supplementary Figure 1: Quality control of recombinant PL25_M23.** **(a)** SDS-PAGE under reducing conditions, Coomassie-stained. M, protein marker; BSA, 2.00 µg bovine serum albumin as a loading reference; R, purified PL25_M23. The protein migrates as a single dominant band near the calculated mass of 36.5 kDa, and purity was assessed as ≥ 90 %. **(b)** Anti-His immunoblot of the same preparation under reducing conditions, probed with a mouse anti-His monoclonal antibody. M, protein marker; R, purified PL25_M23. The major immunoreactive band corresponds to the full-length tagged protein; two faint bands below 20 kDa are also anti-His reactive and, since the tag is N-terminal, are consistent with a minor population of N-terminal fragments. Both panels are from the manufacturer's certificate of analysis for lot U054ZZPXG0-4/P18LA001.

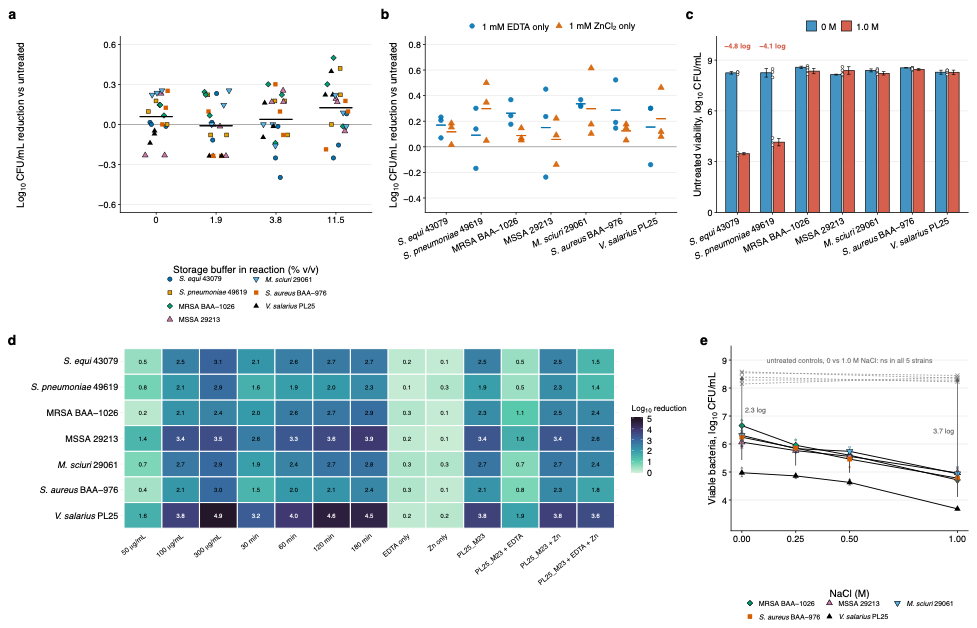

**Supplementary Figure 2: Assay controls for the PL25_M23 antibacterial experiments.** (a) Storage-buffer control: bacteria receiving the PL25_M23-matched volume of protein storage buffer (50 mM Tris-HCl, 150 mM NaCl, 10 % glycerol, pH 8.0) without enzyme, at 0, 1.9, 3.8 and 11.5 % (v/v) of the 200-µL reaction, corresponding to 0, 50, 100 and 300 µg/mL PL25_M23. (b) Additive-only controls: 1 mM EDTA or 1 mM ZnCl2 without enzyme. (c) Untreated viability in PBS with 0 or 1.0 M additional NaCl, showing the losses that led to the exclusion of *S. equi* subsp. *zooepidemicus* ATCC 43079 and *S. pneumoniae* ATCC 49619 from the salt-response analysis; bars are means ± SD with individual replicates shown. (d) Mean log_10_ CFU/mL reduction for every strain and condition tested. (e) The salt-response experiment as raw viability, the data underlying Figure 7b: filled symbols and solid lines, PL25_M23-treated; grey crosses and dashed lines, the untreated controls, which were run at 0 and 1.0 M only; arrows mark the mean separation between the two arms. Untreated viability did not differ between 0 and 1.0 M added NaCl in any retained strain, which is why the 0-M control serves as the reference for all four salt levels in Figure 7b. Panels a–c show Δlog_10_ CFU/mL relative to the matched untreated control, with points as individual biological replicates and horizontal bars as means (n = 3 per strain and condition). All values plotted are listed in Supplementary Data S1.

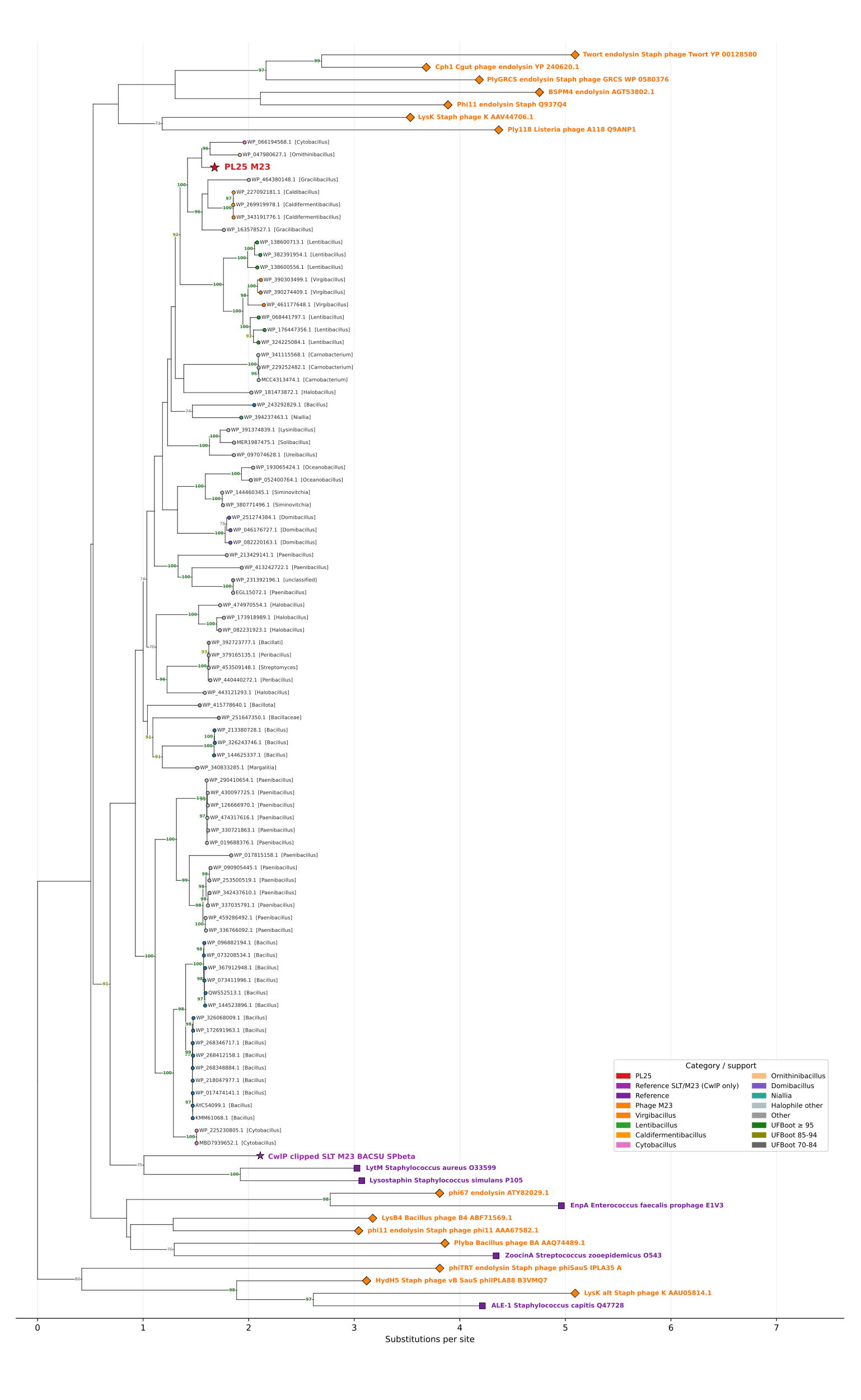

**Supplementary Figure 3: Maximum-likelihood phylogeny of 101 M23-family hydrolases, full topology.** The complete tree underlying Figure 2, shown without collapsing. Sequences comprise PL25_M23 (red star), characterized reference M23 enzymes, phage-associated M23 lysins, and M23-domain proteins recovered from halophilic and halotolerant genera. Sequences were aligned with MAFFT v7.526 (--auto), trimmed with trimAl v1.5.rev1 (-gappyout; 300 retained amino-acid positions), and the tree inferred with IQ-TREE v3.1.1 under the Q.pfam+G4 model with 1,000 ultrafast bootstrap replicates. The tree is midpoint-rooted and branch lengths are substitutions per site. Node labels show ultrafast bootstrap (UFBoot) support and are coloured by support class: green ≥ 95, olive 85–94, grey 70–84; nodes with UFBoot < 70 are unlabelled. Tip symbols and colours indicate taxonomic or functional category as shown in the legend. Bacillus subtilis CwlT was excluded from this phylogeny because its C-terminal peptidase belongs to the NlpC/P60 family (Pfam PF00877) rather than to Peptidase_M23 (PF01551); CwlP is retained as the only published SLT + M23 architectural reference.

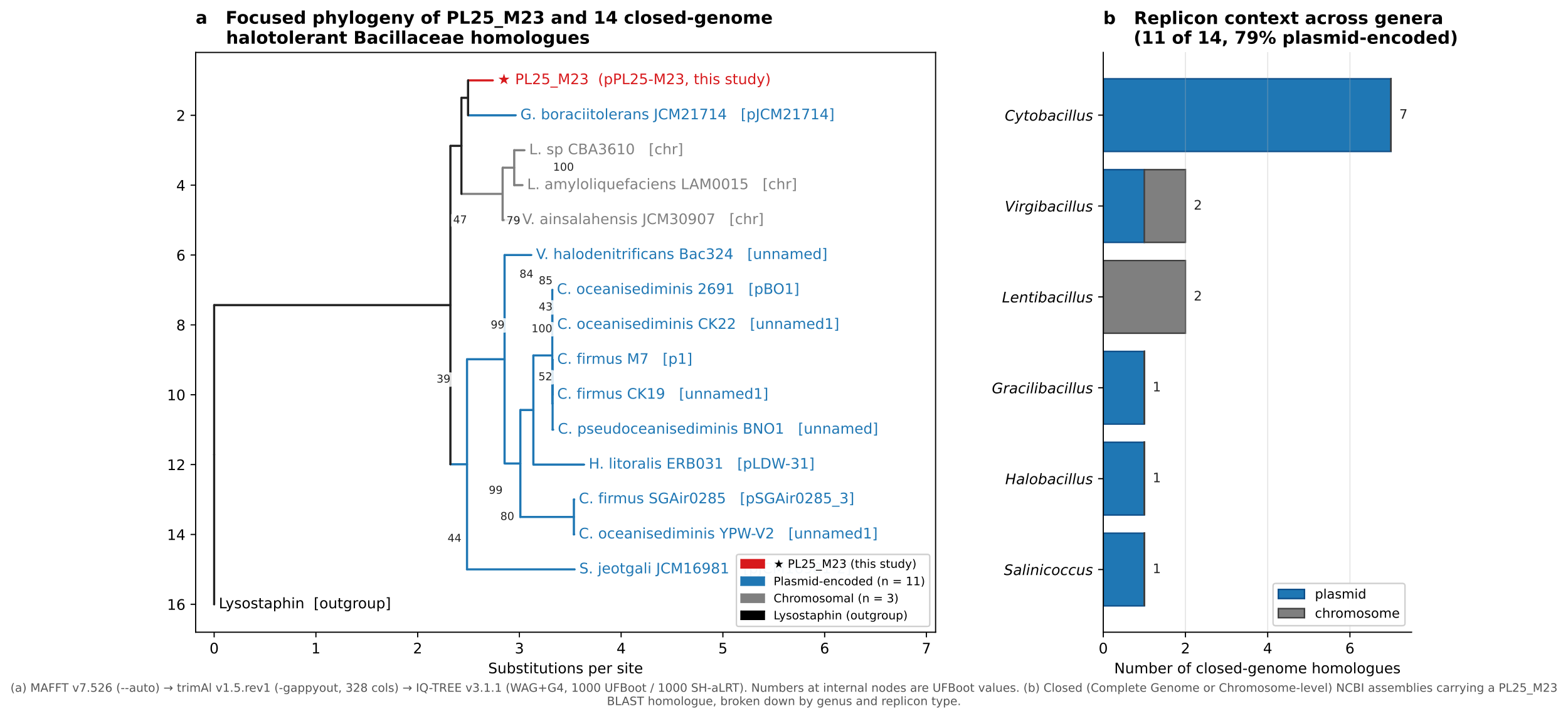

### **Supplementary Figure 4**: **Focused phylogeny and replicon context of the 14 closed-genome PL25_M23 homologues.** (a) Maximum-likelihood phylogeny of PL25_M23, the 14 PL25_M23 homologues encoded in closed NCBI assemblies from the predefined halophilic and halotolerant genus set, and lysostaphin (UniProt P10547) as outgroup, on which the tree is rooted. Full-length protein sequences were aligned with MAFFT v7.526 (--auto, single-threaded), trimmed with trimAl v1.5.rev1 (-gappyout; 328 retained columns), and the tree inferred with IQ-TREE v3.1.1 under WAG+G4, selected by ModelFinder under BIC, with 1,000 ultrafast bootstrap and 1,000 SH-aLRT replicates. Numbers at internal nodes are UFBoot values and branch lengths are substitutions per site. Tip colour indicates replicon context: red, PL25_M23 (this study); blue, plasmid-encoded (n = 11); grey, chromosomally encoded (n = 3); black, outgroup. Plasmid names are given in brackets, and [chr] denotes a chromosomal locus. Replicon assignment was read from the title of the RefSeq nucleotide record linked through the Identical Protein Groups entry for each hit. (b) The same 14 homologues, broken down by genus and replicon type. Accessions, assemblies and BLASTp statistics for every homologue are listed in Supplementary Table S4.
